# Tauopathy severely disrupts homeostatic set-points in emergent neural dynamics but not in the activity of individual neurons

**DOI:** 10.1101/2023.09.01.555947

**Authors:** James N. McGregor, Clayton A. Farris, Sahara Ensley, Aidan Schneider, Chao Wang, Yuqi Liu, Jianhong Tu, Halla Elmore, Keenan D. Ronayne, Ralf Wessel, Eva L. Dyer, Kiran Bhaskaran-Nair, David M. Holtzman, Keith B. Hengen

**Affiliations:** Department of Biology, Washington University in Saint Louis, St. Louis, MO, USA; Department of Physics, Washington University in Saint Louis, St. Louis, MO, USA; Department of Neurology, Hope Center for Neurological Disorders, Knight Alzheimer’s Disease Research Center, Washington University in Saint Louis, St. Louis, MO, USA; Institute for Brain Science and Disease, Chongqing Medical University, 400016, Chongqing, China; Coulter Department of Biomedical Engineering, Georgia Institute of Technology, Atlanta, GA, USA

**Keywords:** Neurodegeneration, criticality, homeostasis, tau, sleep, neurophysiology, behavior, hippocampus

## Abstract

The homeostatic regulation of neuronal activity is essential for robust computation; key set-points, such as firing rate, are actively stabilized to compensate for perturbations. From this perspective, the disruption of brain function central to neurodegenerative disease should reflect impairments of computationally essential set-points. Despite connecting neurodegeneration to functional outcomes, the impact of disease on set-points in neuronal activity is unknown. Here we present a comprehensive, theory-driven investigation of the effects of tau-mediated neurodegeneration on homeostatic set-points in neuronal activity. In a mouse model of tauopathy, we examine 27,000 hours of hippocampal recordings during free behavior throughout disease progression. Contrary to our initial hypothesis that tauopathy would impact set-points in spike rate and variance, we found that cell-level set-points are resilient to even the latest stages of disease. Instead, we find that tauopathy disrupts neuronal activity at the network-level, which we quantify using both pairwise measures of neuron interactions as well as measurement of the network’s nearness to *criticality*, an ideal computational regime that is known to be a homeostatic set-point. We find that shifts in network criticality 1) track with symptoms, 2) predict underlying anatomical and molecular pathology, 3) occur in a sleep/wake dependent manner, and 4) can be used to reliably classify an animal’s genotype. Our data suggest that the critical set-point is intact, but that homeostatic machinery is progressively incapable of stabilizing hippocampal networks, particularly during waking. This work illustrates how neurodegenerative processes can impact the computational capacity of neurobiological systems, and suggest an important connection between molecular pathology, circuit function, and animal behavior.

## INTRODUCTION

The brain’s ability to maintain robust function across a lifetime of experience and change is a remarkable feat and central to cognition and behavior. Homeostatic mechanisms, which stabilize neuronal activity in the face of challenges such as learning and development (Abbott and Nelson, 2000), are understood to be necessary for normal brain function (Turrigiano and Nelson, 2004; Turrigiano 2012). Rationally, there is a selective pressure for set-points maintained by homeostatic mechanisms to be computationally optimal - this is supported by a combination of theory, modeling, and biological data (Miller and MacKay, 1994; Abbott and Nelson, 2000; Turrigiano and Nelson, 2004; Ma et al., 2019; O’Byrne and Jerbi, 2022).

However, there are perturbations for which the brain is unable to compensate - principal amongst these is neurodegenerative disease (NDD), which progressively undermines brain function. Through this lens, we reconsider NDD as a failure of homeostatic mechanisms (Styr and Slutsky, 2018; 2019). Considered from this perspective, neurologic symptoms emerge if and when the brain is unable to compensate for underlying change. This concept shapes the trajectory of NDDs: in animal models of amyotrophic lateral sclerosis, respiratory capacity is unimpaired until more than 70% of phrenic motor neurons have died (Seven et al., 2018). Similarly, in Parkinson’s Disease, symptoms emerge only after significant cell death (Cheng et al., 2010). While respiratory and motor symptoms are clear indicators of underlying damage, it is unclear which aspects of brain activity are affected by disease. Explicitly identifying the disruption in computationally-relevant set-points comprises a direct, dynamical mechanism by which to explain diminished brain function.

Capturing the consistent differences between neuronal activity in healthy and NDD-affected brains has proven challenging. This is in part due to the diversity of patterns and dynamics expected under normal conditions (Buzsáki and Mizuseki, 2014; Hengen et al., 2016; Marder, 2012; Eban-Rothschild et al., 2018; Liberti et al., 2016), but also due to the difficulty of measuring neural activity across the full spectrum of free behavior throughout the timescale of disease onset and progression. Due to these limitations, most investigations into the impacts of NDD on neuronal function have focused on acute measurements of first-order features such as excitability and firing rate in relatively simple settings and tasks. Even in this context, results are mixed and conflicting (Busche et al., 2008; 2012; Crimins et al., 2011; Klee et al., 2020; Hatch et al., 2017; Menkes-Caspi et al., 2015; Booth et al., 2016; Kazim et al., 2021). Summarily, the aspects of neuronal activity that most directly and consistently predict cognitive decline in NDDs remains an open question (Palop and Mucke 2010; Huang & Mucke 2012). Insight into the dynamical consequences of NDDs may be gained by considering the impact of disease on established homeostatic set-points in neural activity believed to support biological computation.

There are three levels at which neuronal activity is known to exhibit robust homeostatic control in vivo: 1) neuron-level setpoints such as firing rate (Hengen et al., 2013; Keck et al., 2013; Hengen et al., 2016), 2) second-order correlations in the local microcircuit (Wu et al., 2020), and 3) higher-order population dynamics (Ma et al., 2019). Because single-neuron firing rate homeostasis is believed to arise, at least in part, from a suite of cell-autonomous mechanisms (Turrigiano et al., 1998; Lambo and Turrigiano, 2013; Shepard et al., 2006 Desai et al., 1999), it is reasonable to hypothesize that firing rate homeostasis would be impacted by misfolded intracellular tau. Dysregulation of firing rates would then be expected to have downstream impacts on neuronal interactions.

To directly address the question of which (if any) homeostatic set-points may be a locus of NDD’s impact on brain activity, we evaluated set-points in neuronal activity in continuous hippocampal single unit recordings spanning > 500 d of life in mice overexpressing a human tau transgene (Shi et al., 2017). Across 27,000 h of recordings, we evaluated the impact of age, genotype, cell type, and sleep/wake state on set-points at the level of the neuron, the microcircuit, and the population computational regime. We report that, contrary to our initial hypotheses, neuron-level set-points of firing rate and variance are unaffected by tauopathy. In contrast, shifts in the network’s computational regime are dramatically undermined by disease. We quantify this by measuring the population’s nearness to *criticality*, a nonequilibrium regime of population dynamics that a) maximizes computational capacity, such as complex problem solving, information storage/transmission, entropy, and dynamic range (Shew and Plenz, 2013; Cocchi et al., 2017; Habibollahi et al., 2022; Cramer et al., 2020), and b) serves as a homeostatic set-point in wildtype (WT) animals (Ma et al., 2019). Disruption of criticality coincides with the onset of disease and deteriorates progressively. Such shifts predict an animal’s genotype, co-vary with underlying molecular pathology, and occur in a state-dependent fashion coincident with the disruption of sleep, one of the earliest behavioral signatures of NDD.

## RESULTS

We hypothesized that NDD must progressively disrupt features of neuronal activity that are normally subject to homeostatic constraint in healthy animals. To test this, we selected the P301S/E4 (TE4) mouse, which overexpresses 1N4R human tau containing the P301S mutation (which causes frontotemporal dementia), on a human apolipoprotein E4 (APOE4) background (Shi et al., 2017; Wang et al., 2021). TE4 mice accumulate insoluble, phosphorylated tau protein by 9 months of age, and subsequently exhibit remarkable hippocampal atrophy (Shi et al., 2017; Wang et al., 2021). We conducted long-term (24 h - 189 d, mean = 38 +/- 6.6 d), continuous recordings of extracellular single unit spiking in CA1 hippocampus prior to and throughout disease progression (Figure 1A-C; 179,478 total single units). Well isolated single units were selected based on waveform properties, spiking statistics, and signal to noise ratio (Supplemental Figure 1; Hengen et al., 2016; Chung et al., 2017; Buccino et al., 2020). Amongst all single units, two cell-types were identified based on well-established criteria: principal cells (excitatory), and fast-spiking inhibitory interneurons (Skaggs et al., 1996; Frank et al., 2001). We report data from 43 animals, comprising 13 TE4, 23 WT, and 7 E4 controls. Consistent with prior reports (Shi et al., 2017, Wang et al., 2021), WT and E4 animals did not display significant differences in neuronal activity and were grouped together as WT/E4 controls (Supplemental Figure 2). Recordings spanned ages between postnatal day 36 (P36) and P533, and were terminated at key points for comparison of recent neuronal dynamics with postmortem molecular and anatomical features of disease progression. Coarse binning by age was guided by prior description of the TE4 mouse, where young (< 3 months), middle (3 - 9 months), and old (> 9 months) capture predisease, onset and early disease, and late disease, respectively (Shi et al., 2017; Wang et al., 2021).

**Figure 1.**
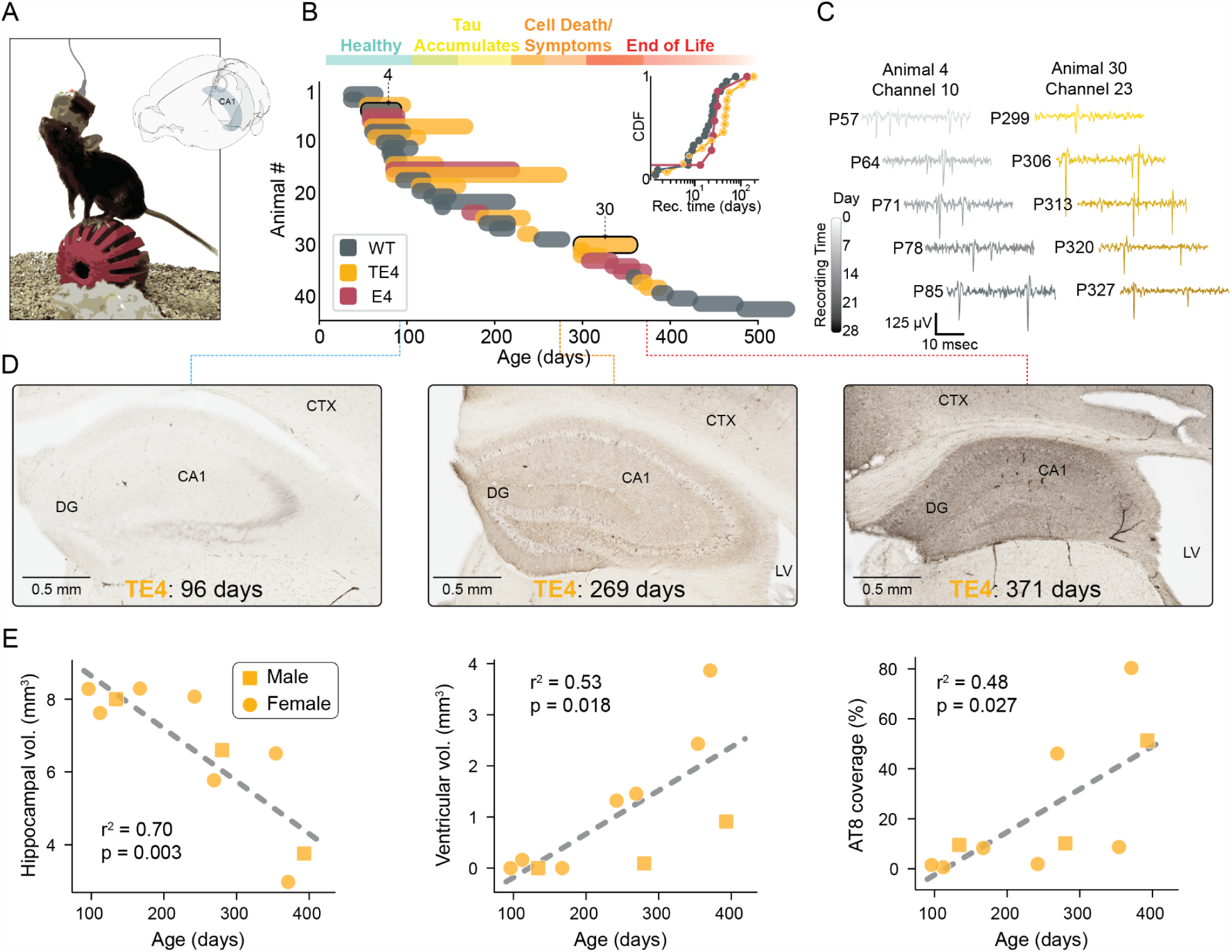
Long-term, continuous recordings of hippocampal neurons in freely behaving WT and tauopathy animals. (A) Experimental approach. Custom electrode arrays were implanted in the CA1 subfield of the hippocampus of mice between the ages of ∼ P50 and P500. Continuous recordings were conducted in the homecage. (B) Recording timelines of 43 animals. Individual recordings were maintained for between 24 h and 189 d (mean = 38 +/- 6.6 d). Each animal’s recording is represented by a colored horizontal bubble; bubbles numbered 4 and 30 correspond to data in the next panel. Gray = WT, red = E4 controls, and orange = TE4 (tauopathy). The typical timecourse of TE4 disease progression is presented along the top. The inset cumulative density plot shows the distribution of recording durations by genotype. (C) Examples of raw data from the same wire over 28 d in a young WT animal (left, gray) and old TE4 animal (right, orange). High signal-to-noise ratio extracellular action potentials are evident through the span of each animal’s data. Animal numbers correspond to those labeled in B. (D) Representative histological images of TE4 animals stained for p-tau (brown) at three timepoints: young (96 d), mid life (269 days), and old (371 d). LV: lateral ventricle, CTX: isocortex, DG: dentate gyrus, CA1: hippocampal subfield. (E) In TE4 animals, hippocampal volume (left), ventricular volume (center), and AT8 coverage correlate with age. Male/female animals are denoted by squares and circles respectively.

To track brain state-specific set-points in neural activity, broadband neural data was synchronized with high-framerate video of animal behavior. Expert human sleep scorers manually labeled 1,440 hours of data (from n = 30 animals) using local field potentials (0.1 - 60 Hz spectral power) and movement data (Xu et al., 2022; Parks et al., 2023). Three broad behavioral states were identified: rapid eye movement (REM) sleep, non-REM sleep (NREM) and wake. Manually labeled data were used to train an XGBoost (extreme gradient boosted decision tree; Chen et al., 2016) tasked with determining behavioral state based purely on neural data (Supplemental Figure 3). The XGBoost achieved a balanced accuracy of 82.7% when tested against data from withheld animals and successfully identified all three behavioral states, including microarousals following REM bouts, consistent with expert human scoring. The output of XGBoost, when applied to > 26,000 h of data from all animals in all conditions, is denoted with a “p” to indicate “probable” (e.g., NREM_p_).

Robust single unit activity was evident throughout all recordings at all stages of life across both genotypes (TE4 and WT/E4; Figure 1C). Animals were sacrificed intermittently between P90 and P550; TE4 animals exhibited progressive neurodegeneration and tau accumulation after P200 (Figure 1D,E).

### Single neuron firing rates are robust to age and disease status

Homeostatic set-points in the firing rates of individual neurons are continuously regulated on the timecourse of hours to days (Hengen et al., 2016; Pacheco et al., 2021; Keck et al., 2013). We reasoned that the accumulation of intracellular tau aggregates might undermine the ability of neurons to maintain firing rate set-points, which vary by cell type and brain state (Csicsvari et al., 1998; Hengen et al., 2013, 2016; Xu et al., 2022). To address this, we considered the central tendency of all recorded single units as a function of cell type and arousal state.

Mean firing rates of individual neurons were lognormally distributed as expected (Buszaki and Mizuseki, 2014; Mizuseki et al., 2013), and ensembles of both excitatory and inhibitory neurons exhibited stable rates across multiple weeks, consistent with prior descriptions of firing rate set-points (Hengen et al., 2013, 2016). This was evident in both young and old animals as well as WT/E4 and TE4 (Figure 2A-C). As expected, pFS neurons exhibited higher mean rates than principal neurons (7.91 +/- 1.19 versus 2.09 +/- 0.22 Hz, respectively, p < 0.0001, t-test), each of whose firing rates were lower in NREM_p_ than wake_p_ (both cell types p < 0.0001, linear mixed effects regression: log_firing_rate ∼ age_group * genotype * state * cell_type + animal, where animal is included as a random effect). pFS interneuron rates were lowest during NREM_p_ and highest during REM_p_ (p < 0.0001 for each pairwise comparison), while principal neuron rates were lowest during REM_p_ and highest during wake (NREM_p_ versus REM_p_: p = 0.019, other pairwise contrasts p < 0.0001)(see Table 1).

**Figure 2.**
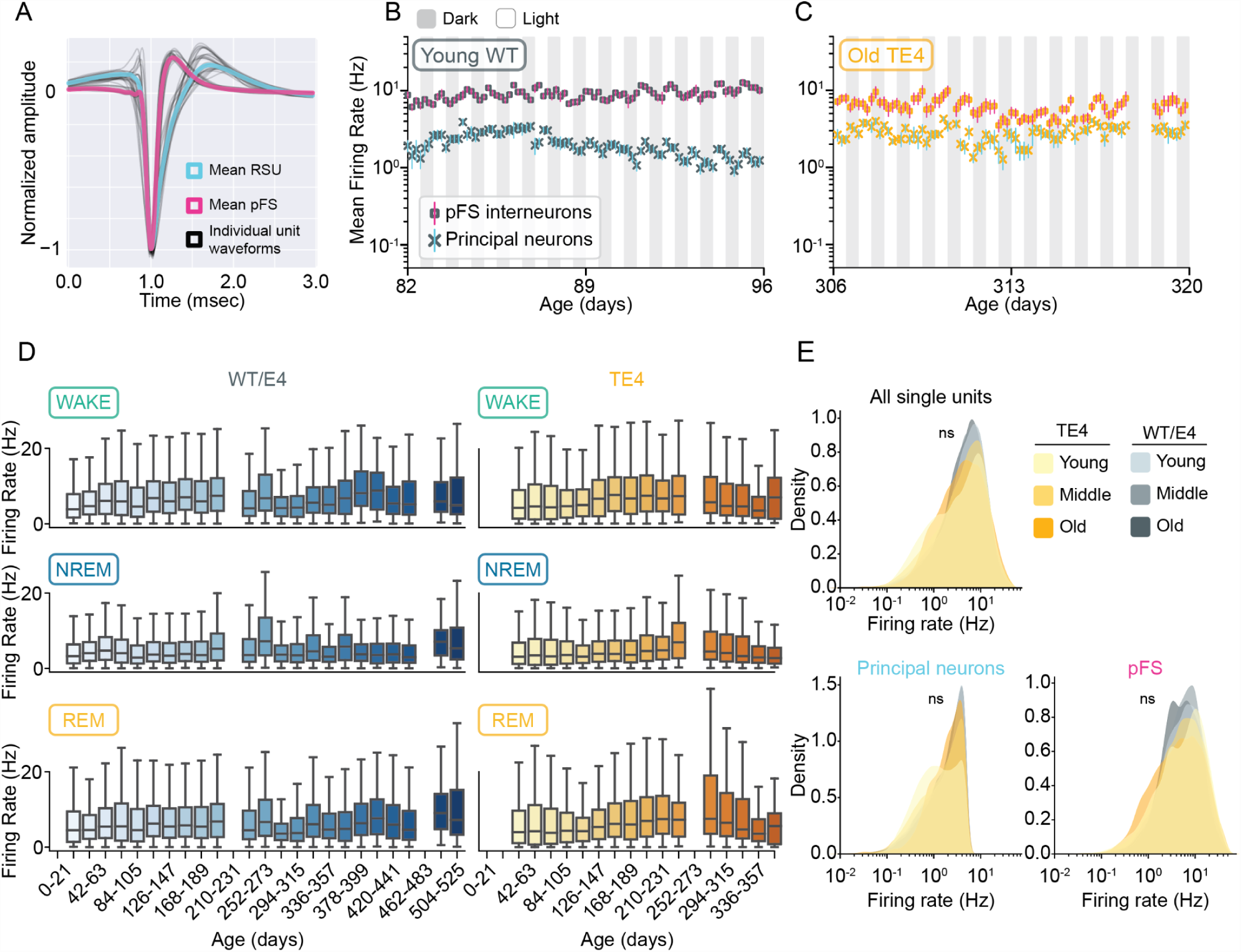
The mean firing rate of single neurons in CA1 is resilient to tauopathy and old age. (A) Single units were separated into regular spiking units (RSU: blue) and putative fast spiking inhibitory interneurons (pFS: pink). A subset of RSUs were further identified as principal cells. (B) Example of stable ensemble mean firing rate over 14 d of continuous recording in a young WT animal. Xs with blue error denote the mean principal cell rate. Squares with pink error denote the mean pFS rate. (C) Same as B but ensemble data were collected in an old TE4 animal. (D) The mean firing rate of all well-isolated single units across the lifetime as a function of brain state (rows) and genotype (columns). (E) Kernel density estimates of single unit mean firing rates as a function of age group (young, middle, and old) and genotype (orange = TE4, gray = WT/E4). The top panel shows all single units, while the bottom panels show the distributions of firing rates for principal neurons (left) and pFS neurons (right). For pairwise statistical contrasts, see Tables 2 - 4.

**Table 1.** Single neuron mean firing rates as a function of age, genotype, cell type, and arousal state. Each well isolated single unit was evaluated as a function of NREM sleep, REM sleep, and wake. The mean firing rate (Hz) was calculated by state for each neuron. Means were then averaged by condition (e.g., genotype). Error is SEM, where n animals is used in the denominator.

| Age | Genotype | Cell Type | NREM | REM | Wake |
| --- | --- | --- | --- | --- | --- |
| Young | WT | Principal | 2.1 ± 0.4 | 2.4 ± 0.6 | 2.5 ± 0.5 |
|  |  | pFS | 6.2 ± 1.9 | 9.6 ± 2.8 | 8.8 ± 2.5 |
|  | TE4 | Principal | 1.4 ± 0.6 | 1.5 ± 0.9 | 1.7 ± 0.7 |
|  |  | pFS | 7.6 ± 3.7 | 10.9 ± 5.5 | 9.1 ± 4.1 |
| Middle | WT | Principal | 1.7 ± 0.3 | 2.3 ± 0.5 | 2.6 ± 0.5 |
|  |  | pFS | 6.5 ± 1.9 | 9.4 ± 2.4 | 9.5 ± 2.3 |
|  | TE4 | Principal | 1.8 ± 0.5 | 2.2 ± 0.8 | 2.3 ± 0.7 |
|  |  | pFS | 6.7 ± 2.8 | 10.2 ± 4.0 | 8.9 ± 3.2 |
| Old | WT | Principal | 2.0 ± 0.5 | 2.3 ± 0.6 | 2.5 ± 0.6 |
|  |  | pFS | 5.6 ± 2.0 | 7.9 ± 2.8 | 7.5 ± 2.4 |
|  | TE4 | Principal | 2.3 ± 0.7 | 3.1 ± 1.0 | 2.5 ± 0.7 |
|  |  | pFS | 7.5 ± 4.3 | 11.9 ± 6.7 | 8.8 ± 4.5 |

To address the impact of tauopathy on firing rate set-points, we tracked state-dependent set-points in pFS and principal neuron rates between P36 and P533 as a function of genotype (Figure 2D; Supplemental Figure 4). To our surprise and contrary to our primary hypothesis, tauopathy had no measurable impact on the mean firing rate of either cell type in any of the three behavioral states. Specifically, when considering firing rates, there was no main effect of age (p = 0.459), no main effect of genotype (p = 0.608), and no significant interaction between the two (p = 0.394; linear mixed effects regression, log_firing_rate ∼ age * genotype * state* cell_type + 1|animal, where animal is included as a random effect). There was a significant 3-way interaction between age, cell type, and genotype, however pairwise comparisons revealed, at most, differences of <1.6 Hz unlikely to be of biological significance (Supplemental Tables 2 - 4). For comparison, sensory deprivation drives a ∼ 60% reduction in firing rate (Hengen et al., 2016).

**Tables 2 - 4.** Statistical comparisons of mean firing rate by cell type, age, and genotype. Single unit mean firing rates were calculated and grouped by cell type, age, and genotype. Due to the log-normal distribution of mean rates, they were log transformed before further analysis. Data were fit using a linear mixed effects model with the formula: log_fr ∼ age_group * genotype * cell_type + (1∣animal) where ’animal’ is included as a random effect to account for repeated measures and inherent variability across different animals. Pairwise comparisons were conducted using the emmeans package. Tables provide contrasts among the levels of the fixed effects and the associated p-values. All p-values were adjusted using Tukey’s method to account for multiple comparisons. Contrast: Comparisons between different conditions. Estimate: Predicted difference between the groups in the given contrast. SE (Standard Error): Measure of variability around the estimated difference, accounting for both fixed and random effects. z-ratio: z-ratio: The estimated difference divided by its standard error, used to test the significance of the contrast. p-value: Probability of observing the given z-ratio under the null hypothesis of no difference.

| Contrast | Estimate | SE | z-ratio | P-value |
| --- | --- | --- | --- | --- |
| te4 mid - wt mid | -0.03376 | 0.04463 | -0.757 | 0.9746 |
| te4 mid - te4 old | 0.04923 | 0.06339 | 0.777 | 0.9715 |
| te4 mid - wt old | 0.01158 | 0.05107 | 0.227 | 0.9999 |
| te4 mid - te4 young | -0.04179 | 0.00518 | -8.075 | <0.0001 |
| te4 mid - wt young | -0.03841 | 0.04468 | -0.860 | 0.9559 |
| wt mid - te4 old | 0.08300 | 0.05591 | 1.484 | 0.6743 |
| wt mid - wt old | 0.04534 | 0.04142 | 1.095 | 0.8837 |
| wt mid - te4 young | -0.00803 | 0.04475 | -0.179 | 1.0000 |
| wt mid - wt young | -0.00465 | 0.00459 | -1.013 | 0.9139 |
| te4 old - wt old | -0.03765 | 0.06117 | -0.616 | 0.9899 |
| te4 old - te4 young | -0.09102 | 0.06347 | -1.434 | 0.7062 |
| te4 old - wt young | -0.08764 | 0.05595 | -1.566 | 0.6209 |
| wt old - te4 young | -0.05337 | 0.05117 | -1.043 | 0.9034 |
| wt old - wt young | -0.04999 | 0.04148 | -1.205 | 0.8345 |
| te4 young - wt young | 0.00338 | 0.04480 | 0.075 | 1.0000 |

| Contrast | Estimate | SE | z-ratio | P-value |
| --- | --- | --- | --- | --- |
| te4 mid - wt mid | -0.02235 | 0.04463 | -0.501 | 0.9962 |
| te4 mid - te4 old | -0.19121 | 0.06344 | -3.014 | 0.0309 |
| te4 mid - wt old | -0.06752 | 0.05107 | -1.322 | 0.7730 |
| te4 mid - te4 young | 0.05824 | 0.00528 | 11.024 | <0.0001 |
| te4 mid - wt young | -0.10453 | 0.04466 | -2.341 | 0.1778 |
| wt mid - te4 old | -0.16886 | 0.05597 | -3.017 | 0.0307 |
| wt mid - wt old | -0.04517 | 0.04143 | -1.090 | 0.8855 |
| wt mid - te4 young | 0.08059 | 0.04477 | 1.800 | 0.4656 |
| wt mid - wt young | -0.08218 | 0.00459 | -17.89 | <0.0001 |
| te4 old - wt old | 0.12369 | 0.06123 | 2.020 | 0.3307 |
| te4 old - te4 young | 0.24944 | 0.06353 | 3.926 | 0.0012 |
| te4 old - wt young | 0.08668 | 0.05599 | 1.548 | 0.6329 |
| wt old - te4 young | 0.12575 | 0.05119 | 2.457 | 0.1373 |
| wt old - wt young | -0.03701 | 0.04146 | -0.893 | 0.9484 |
| te4 young - wt young | -0.16277 | 0.04479 | -3.634 | 0.0038 |

| Contrast | Estimate | SE | z-ratio | P-value |
| --- | --- | --- | --- | --- |
| te4 mid - wt mid | -0.008117 | 0.03047 | -0.266 | 0.9998 |
| te4 mid - te4 old | -0.001963 | 0.04172 | -0.047 | 1.0000 |
| te4 mid - wt old | 0.000483 | 0.03531 | 0.014 | 1.0000 |
| te4 mid - te4 young | -0.004162 | 0.00357 | -1.165 | 0.8534 |
| te4 mid - wt young | -0.031611 | 0.03051 | -1.036 | 0.9058 |
| wt mid - te4 old | 0.006154 | 0.03655 | 0.168 | 1.0000 |
| wt mid - wt old | 0.008600 | 0.02850 | 0.302 | 0.9997 |
| wt mid - te4 young | 0.003955 | 0.03057 | 0.129 | 1.0000 |
| wt mid - wt young | -0.023494 | 0.00397 | -5.924 | <0.0001 |
| te4 old - wt old | 0.002445 | 0.04067 | 0.060 | 1.0000 |
| te4 old - te4 young | -0.002199 | 0.04179 | -0.053 | 1.0000 |
| te4 old - wt young | -0.029648 | 0.03658 | -0.811 | 0.9657 |
| wt old - te4 young | -0.004645 | 0.03539 | -0.131 | 1.0000 |
| wt old - wt young | -0.032094 | 0.02855 | -1.124 | 0.8716 |
| te4 young - wt young | -0.027449 | 0.03061 | -0.897 | 0.9474 |

While well-isolated single units were separated into pFS and principal neurons, there were a number of putative excitatory cells whose firing rate precluded them from the principal class (5 Hz cap; Skaggs et al., 1996; Csicsvari et al., 1999; Frank et al., 2001; Mizuseki et al., 2012; Booth et al., 2016). This raises the possibility that increases in principal neuron firing rate are obscured simply because affected cells are excluded from analyses. To evaluate this, we examined the full distribution of mean firing rates of *all* well-isolated single units (Fig. 2E). Across all neurons, there was not a significant main effect of genotype (p = 0.655, linear mixed effects regression). Over 15 pairwise comparisons across age groups and genotypes, only one significant contrast was evident (WT/E4 young vs WT/E4 middle age, p < 0.0001), although the mean difference was 1.05 Hz, or 16 % of the mean (Table 4).

To account for the possibility of overfitting a large dataset, we deployed two complementary statistical approaches to testing whether mean firing rates differed by genotype and age. First, we used a hierarchical bootstrap test which considers a balanced subset of 10,000 neurons (McGregor et al., 2022; Saravanan et al., 2020; Efron, 1982). This yielded no significant differences between any combination of ages and/or genotypes (0.20 < P_boot_ < 0.71; Supplemental Figure 4C). Next, to ask whether an effect of genotype would be detectable in shorter intervals, we used a bootstrapped linear mixed model. Within the three age ranges, we randomly selected three 2 h bins of data from each animal and tested whether the mean firing rates of WT/E4 and TE4 neurons were different. We then repeated this process 100 times to generate a range of p-values expected in acute observations of neuronal activity. In young animals, 2/100 tests yielded a p-value of < 0.05. In middle-aged animals, 1/100 tests yielded a p-value of < 0.05. In old animals, 5/100 tests yielded a p-value of < 0.05 (Supplemental Figure 4D).

Taken together, these data point to a surprising conclusion: mean firing rate, a key parameter subject to homeostatic control in the CNS, is not substantially impacted by either tauopathy or age. However, our analyses are blind to neurons that have died or fallen silent. Given that post mortem histology revealed hippocampal atrophy (Figure 1D,E), which of other studies is concomitant with significant neuronal loss (Shi et al., 2017; Wang et al., 2021; Chen et al., 2023), these data suggest that, until a neuron is no longer detectable by our methods, the mechanisms supporting homeostatic regulation of firing rate are intact.

### Spike time variance is unaffected by tauopathy

Despite the apparent immutability of firing rate, we reasoned that tau’s impact on neuronal activity may still be at the level of the single neuron. Downstream of firing rate, there are homeostatic set-points in single neuron activity that have powerful implications for biological computation and thus brain function. Key amongst these is the irregularity of spike timing, which can be quantified by the coefficient of variation (CV) of a neuron’s interspike interval (ISI) distribution (Hengen et al., 2013). In the same datasets presenting stable rates, we examined CV as a function of age, genotype, cell type, and sleep/wake state.

Consistent with prior reports of in vivo neuronal activity (Hengen et al., 2013; Urbain et al., 2006), the mean CV of ISI of each cell type was slightly super-poisson (pFS = 1.37 +/- 0.07, principal = 1.53 +/- 0.07, pairwise contrast p < 0.0001, linear mixed effects regression). Similar to firing rate data, the mean CV of ensembles of both excitatory and inhibitory neurons was stable over multiple weeks in both young and old animals as well as WT/E4 and TE4 (Figure 3A,B; Hengen et al., 2013). Each cell type exhibited state dependent effects that did not parallel those observed in firing rates. Specifically, in pFS neurons, CV was significantly higher during wake_p_ (1.47 +/- 0.09) than either sleep state (NREM_p_: 1.42 +/- 0.05, REM_p_: 1.44 +/- 0.1, p < 0.0001 for each comparison to wake, linear mixed effects). The sleep states were not different from one another (NREM_p_ vs REM_p_, p = 0.67). Principal neuron CV was significantly higher during wake_p_ than REM_p_ sleep (1.71 +/- 0.1 vs 1.56 +/- 0.09, p < 0.0001), which was significantly higher than NREM_p_ sleep (1.43 +/- 0.05, p < 0.0001). These state-dependent changes in CV of ISI were consistent with prior literature (Evarts et al., 1962; Hengen et al., 2016). Taken together, the reliability of these values indicate state-dependent set-points in spike time variance (Table 5).

**Figure 3.**
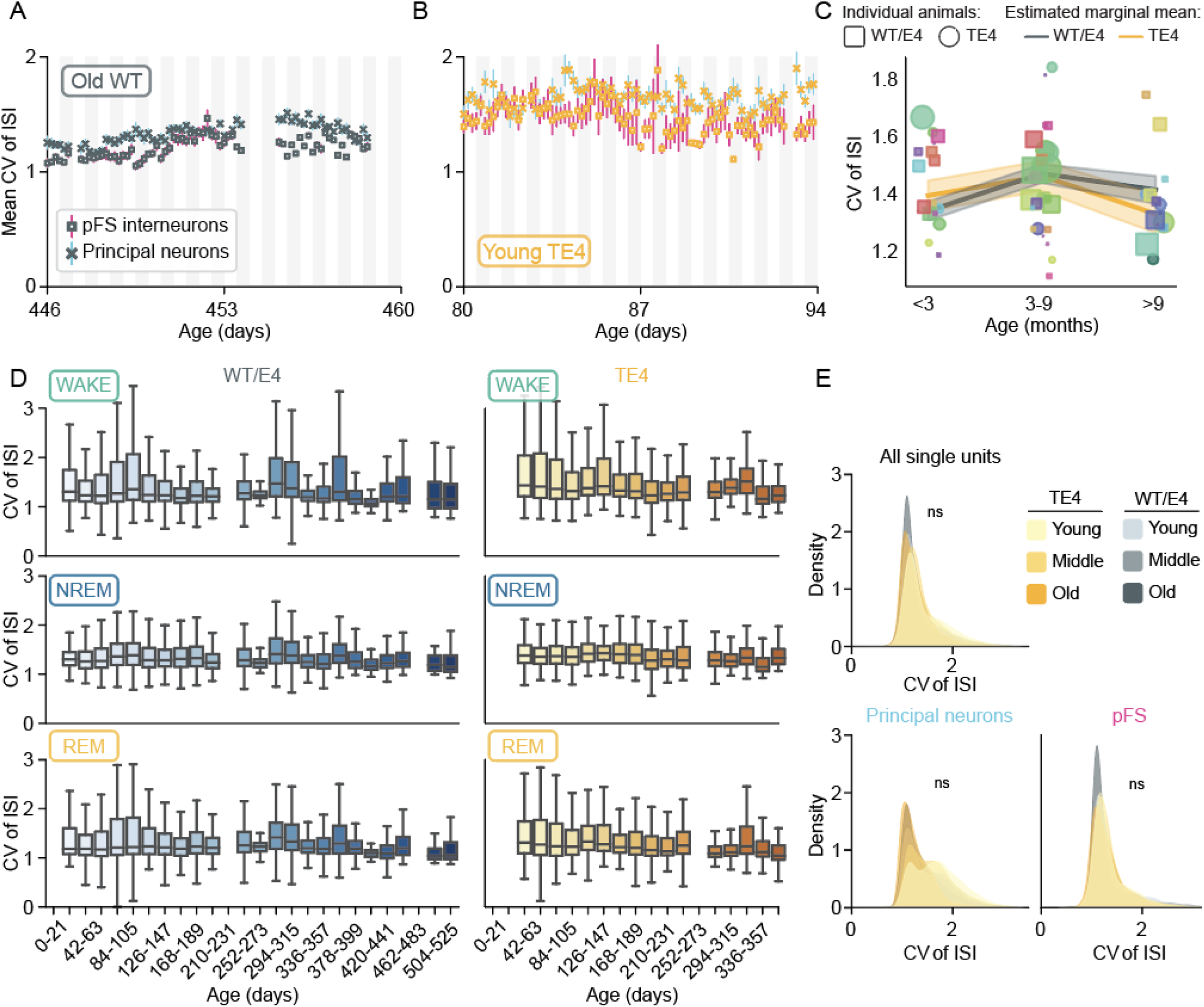
Set-points in spike time variance are robust to tauopathy and old age. (A) Example of mean coefficient of variation (CV) of interspike intervals (ISIs) in an ensemble of principal neurons (Xs with blue error) and pFS interneurons (squares with pink error) over 14 d of continuous recording in an old WT animal. (B) Same as A, but example is from an ensemble recorded in a young TE4 animal. (C) Estimated marginal means of CV of ISI of all single units were extracted from the linear mixed model that accounts for individual animals and variable durations of observation (CV_of_ISI ∼ genotype * age_group * cell_type + (1|animal) where animal is included as a random effect). Estimated marginal means are plotted as lines across age group and colored by genotype (error is model SE). Scatter points show the means of individual animals, indicated by color. Squares are WT/E4 animals, circles are TE4 animals. (D) The mean CV of ISIs of all well-isolated single units across the lifetime as a function of brain state (rows) and genotype (columns). (E) Kernel density estimates of single unit ISI CVs as a function of age group (young, middle, and old) and genotype (orange = TE4, gray = WT/E4). The top panel shows all single units, while the bottom panels show the distributions of ISI CVs for principal neurons (left) and pFS neurons (right). For pairwise statistical contrasts, see Tables 6 - 8.

**Table 5.**
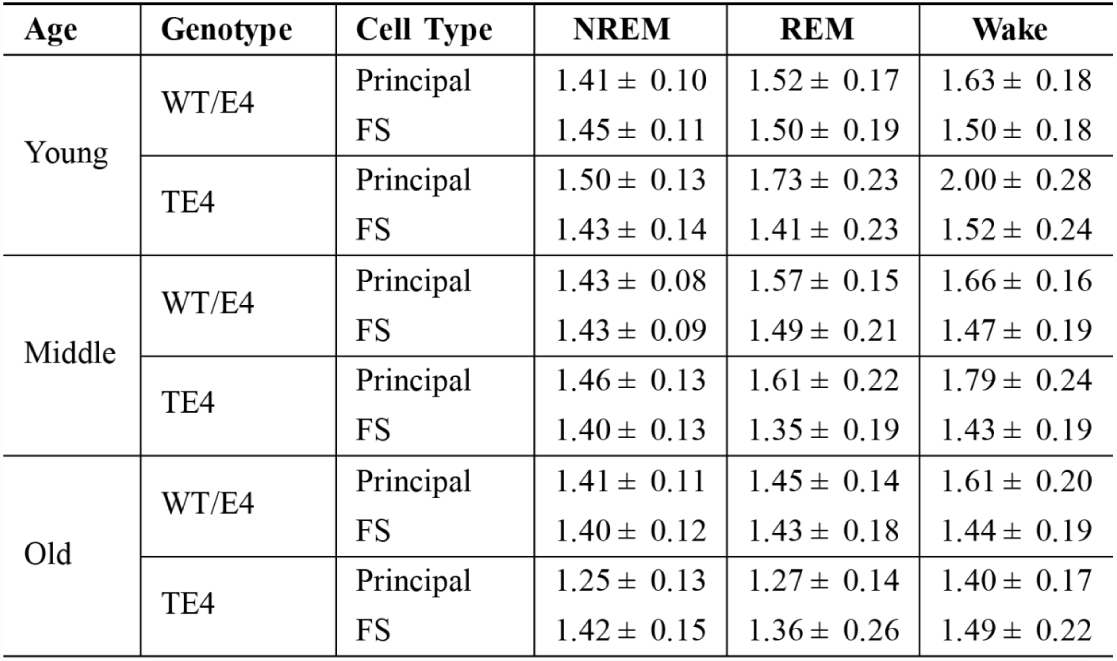
Mean single neuron ISI CV as a function of age, genotype, cell type, and arousal state. The coefficient of variation (CV) of the interspike interval (ISI) of each well isolated single unit was evaluated as a function of NREM sleep, REM sleep, and wake. The mean CV was then calculated by condition (e.g., genotype). Error is SEM, where n animals is used in the denominator.

**Tables 6 - 8.** Statistical comparisons of single unit CV of ISI by cell type, age, genotype and brain state. The coefficient of variation (CV) of single neuron interspike intervals (ISIs) were calculated for each brain state. The data were fit using a linear mixed effects model with the formula: cv ∼ age_group * genotype * state * cell_type + (1∣animal) where ’animal’ is included as a random effect to account for repeated measures and inherent variability across different animals. Pairwise comparisons were conducted using the emmeans package. Tables provide contrasts among the levels of the fixed effects and the associated p-values. All p-values were adjusted using Tukey’s method to account for multiple comparisons. Estimate: Predicted difference between the groups in the given contrast. SE (Standard Error): Measure of variability around the estimated difference, accounting for both fixed and random effects. z-ratio: z-ratio: The estimated difference divided by its standard error, used to test the significance of the contrast. p-value: Probability of observing the given z-ratio under the null hypothesis of no difference.

| Contrast | Estimate | SE | z-ratio | P-value |
| --- | --- | --- | --- | --- |
| te4 mid - wt mid | -0.05234 | 0.04974 | -1.052 | 0.9000 |
| te4 mid - te4 old | 0.02660 | 0.07062 | 0.377 | 0.9990 |
| te4 mid - wt old | -0.04288 | 0.05689 | -0.754 | 0.9750 |
| te4 mid - te4 young | 0.00406 | 0.00569 | 0.713 | 0.9804 |
| te4 mid - wt young | -0.00510 | 0.04979 | -0.102 | 1.0000 |
| wt mid - te4 old | 0.07894 | 0.06228 | 1.268 | 0.8028 |
| wt mid - wt old | 0.00946 | 0.04612 | 0.205 | 1.0000 |
| wt mid - te4 young | 0.05639 | 0.04986 | 1.131 | 0.8685 |
| wt mid - wt young | 0.04724 | 0.00502 | 9.407 | <.0001 |
| te4 old - wt old | -0.06948 | 0.06813 | -1.020 | 0.9115 |
| te4 old - te4 young | -0.02255 | 0.07071 | -0.319 | 0.9996 |
| te4 old - wt young | -0.03170 | 0.06232 | -0.509 | 0.9959 |
| wt old - te4 young | 0.04693 | 0.05699 | 0.824 | 0.9633 |
| wt old - wt young | 0.03778 | 0.04618 | 0.818 | 0.9643 |
| te4 young - wt young | -0.00916 | 0.04991 | -0.183 | 1.0000 |

| Contrast | Estimate | SE | z-ratio | P-value |
| --- | --- | --- | --- | --- |
| te4 mid - wt mid | 0.04930 | 0.04974 | 0.991 | 0.9210 |
| te4 mid - te4 old | 0.31079 | 0.07068 | 4.397 | 0.0002 |
| te4 mid - wt old | 0.12185 | 0.05690 | 2.142 | 0.2658 |
| te4 mid - te4 young | -0.01086 | 0.00582 | -1.864 | 0.4243 |
| te4 mid - wt young | 0.14591 | 0.04976 | 2.932 | 0.0395 |
| wt mid - te4 old | 0.26149 | 0.06234 | 4.194 | 0.0004 |
| wt mid - wt old | 0.07256 | 0.04613 | 1.573 | 0.6166 |
| wt mid - te4 young | -0.06016 | 0.04987 | -1.206 | 0.8341 |
| wt mid - wt young | 0.09661 | 0.00504 | 19.162 | <.0001 |
| te4 old - wt old | -0.18894 | 0.06819 | -2.771 | 0.0622 |
| te4 old - te4 young | -0.32165 | 0.07077 | -4.545 | 0.0001 |
| te4 old - wt young | -0.16488 | 0.06236 | -2.644 | 0.0870 |
| wt old - te4 young | -0.13271 | 0.05701 | -2.328 | 0.1829 |
| wt old - wt young | 0.02406 | 0.04616 | 0.521 | 0.9954 |
| te4 young - wt young | 0.15677 | 0.04990 | 3.142 | 0.0208 |

| Contrast | Estimate | SE | z-ratio | P-value |
| --- | --- | --- | --- | --- |
| te4 mid - wt mid | 0.01903 | 0.04964 | 0.383 | 0.9989 |
| te4 mid - te4 old | 0.13386 | 0.07039 | 1.902 | 0.4010 |
| te4 mid - wt old | 0.04463 | 0.05676 | 0.786 | 0.9699 |
| te4 mid - te4 young | 0.02170 | 0.00408 | 5.324 | <.0001 |
| te4 mid - wt young | 0.09460 | 0.04965 | 1.905 | 0.3988 |
| wt mid - te4 old | 0.11482 | 0.06202 | 1.851 | 0.4326 |
| wt mid - wt old | 0.02559 | 0.04598 | 0.557 | 0.9937 |
| wt mid - te4 young | 0.00266 | 0.04971 | 0.054 | 1.0000 |
| wt mid - wt young | 0.07556 | 0.00384 | 19.694 | <.0001 |
| te4 old - wt old | -0.08923 | 0.06785 | -1.315 | 0.7768 |
| te4 old - te4 young | -0.11216 | 0.07043 | -1.592 | 0.6036 |
| te4 old - wt young | -0.03926 | 0.06203 | -0.633 | 0.9886 |
| wt old - te4 young | -0.02293 | 0.05682 | -0.404 | 0.9986 |
| wt old - wt young | 0.04997 | 0.04598 | 1.087 | 0.8869 |
| te4 young - wt young | 0.07290 | 0.04972 | 1.466 | 0.6859 |

In this context, we asked whether tauopathy progressively disrupts set-points in the timing of neuronal spiking. Contrary to our expectations, across all single units (pFS, principal, and other) we observed no meaningful differences in CV as a function of age or genotype (Figure 3C, Table 6-8. Main effect of genotype p = 0.98; main effect of age p < 0.0001: young = 1.36 vs mid-life = 1.46). Similarly, state-specific set-points in CV did not meaningfully change as a function of age or genotype (Figure 3D). Broken up by cell type, the same trend held: there were no meaningful differences either at the level of cell type (Figure 3E) or in the interactions of cell type, age, and behavioral state (Supplemental Figure 5).

Small yet significant differences that emerged when considering the entire dataset were not evident when CV was evaluated using hierarchical bootstrapping (0.17 < P_boot_ < 0.73) or a bootstrapped linear mixed effects regression on shorter intervals of data (0/100 tests comparing WT/E4 and TE4 yielded p < 0.05 in each of the three age groups (Supplemental Figure 5C,D). These results imply that the functional effects of tauopathy are likely to be explained by disruptions in neuronal activity unrelated to the ability of individual neurons to maintain local set-points.

### Ensemble pairwise correlation features are impacted by tauopathy

In addition to constraints in single neuron activity, there are homeostatic set-points at the level of network interactions. The simplest example of a network-level set-point is the average strength of pairwise correlations in an ensemble (Wu et al., 2020). To evaluate this, we binned neuronal activity at 100 msec and calculated the mean pairwise correlation between all pairs of single units in 30 minute windows as described in Wu et al., 2020. We performed hierarchical clustering in the first 30 min window and maintained the row/column assignments of each neuron as we stepped the window forward in 5 min increments (Figure 4A). This approach enabled us to measure three features of the pairwise correlation matrix: 1) the mean pairwise correlation strength, which is subject to homeostatic control (Wu et al., 2020), 2) the stability of the pairwise correlation matrix over time, as measured by the mean normalized L1 distance between matrices separated by 4 h, and 3) the entropy of the correlation matrix (bits per pair), a measure of its complexity.

**Figure 4.**
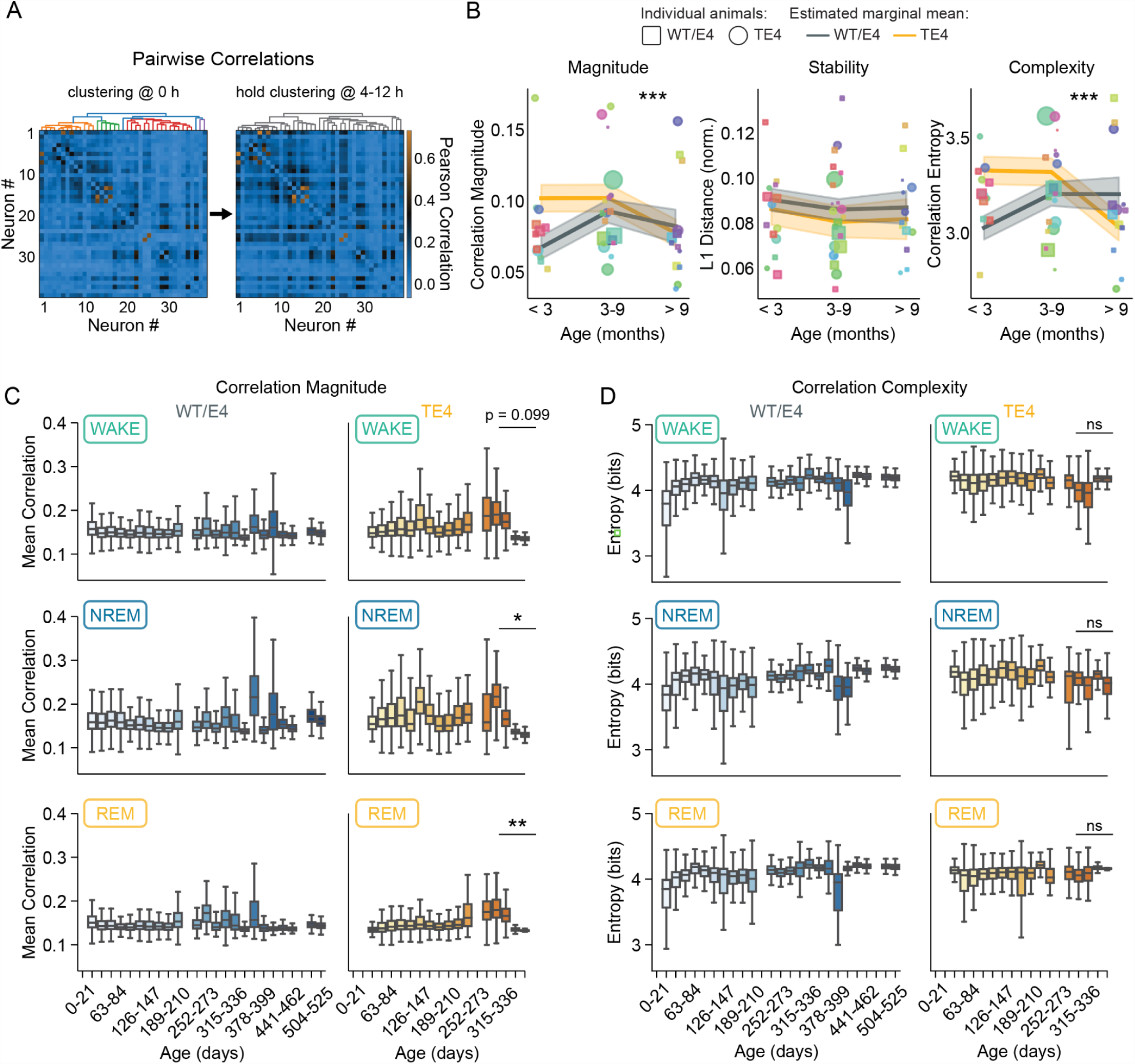
Set-points in ensemble pairwise correlations may be impacted by tauopathy. (A) Example correlation heatmap showing all pairwise comparisons of single units at the beginning (left) and end (right) of a 12 h data block. Neuron ordering is determined by hierarchical clustering at the beginning of the block (colored dendrogram, top left) and maintained throughout the block (gray dendrogram, top right) to allow visualization of the stability of the correlation structure. (B) Across all recordings, three features of pairwise correlations were evaluated. The ensemble mean pairwise correlation magnitude (left), the average stability of the correlation matrix across 4 h intervals (middle), and the complexity (right) of the correlation matrix (entropy: mean bits/pair). Animal means are shown by scatter, where the size of each point is proportional to the amount of data. Fit lines show the estimated marginal means based on linear mixed effects (correlation_feature ∼ age_group * genotype * state * cell_type + (1|animal), where animal is included as a random effect). Asterisks denote pairwise contrasts of old TE4 to young/mid TE4. (C) Mean correlation magnitude across the lifetime as a function of sleep/wake state (rows) and genotype (columns). Asterisks show the pairwise contrast of old TE4 to old WT/E4. (D) Same as E but the mean correlation matrix complexity. *** p < 0.0001, ** p < 0.01, * p < 0.05. For complete pairwise contrasts, see Tables 9-11.

Broadly, there was no main effect of genotype (correlation magnitude p = 0.27; stability p = 0.47; entropy p = 0.36; linear mixed effects), and no instances in which WT/E4 and TE4 pairwise contrasts were significant (Figure 4B). Despite this, there were noteworthy and significant changes in correlation magnitude and complexity between mid-life and late life in TE4 but not WT/E4 animals. Specifically, correlation magnitude decreased between mid (0.10 +/- 0.01) and old ages (0.08 +/- 0.01) in TE4 animals (interaction of age and genotype p <2e-16; TE4 mid versus old p < 0.0001). Similarly, the entropy of the pairwise correlation matrix also decreased in old TE4 animals compared to mid-life (mid-life: 3.32 +/- 0.07 bits/pairs; old: 3.03 +/- 0.07 bits/pairs, p < 0.0001). In contrast, there was no meaningful age or genotype dependent difference in the stability of the pairwise correlation matrix (age by genotype interaction p = 0.934).

Given state-dependent set-points in firing rate and CV, we asked whether a) there are sleep/wake set-points in the pairwise correlation structure, and b) whether these set-points might be sensitive to tauopathy. Linear mixed effects regression revealed a powerful main effect of state in both correlation magnitude and complexity (Figure 4C,D. Magnitude: NREM_p_ > wake_p_ > REM_p_, all comparisons p < 0.0001. Complexity: wake_p_ > NREM_p_ > REM_p_, all comparisons p < 0.0001. Corr_measure ∼ genotype * sleep/wake * age_group + 1|animal, where animal is included as a random effect). This suggests that, independent of age and genotype, state consistently influences the structure of hippocampal pairwise correlations. Further, magnitude significantly differed between old TE4 and old WT animals in each of the two sleep states, but not wake (old TE4 vs old WT/E4: NREM_p_ p = 0.019; REM_p_ p = 0.007, wake_p_ p = 0.099. Figure 4C, Tables 9-11). It is worth noting that the two oldest TE4 animals (> 1y) are characterized by a distinct drop off in correlation magnitude that appears to begin around 7 months of age (Figure 4C). Finally, the stability of the correlation matrix was statistically indistinguishable between sleep_p_-dense and wake_p_-dense blocks in both genotypes across all age groups (Supplemental Figure 6C).

**Tables 9 - 11.**
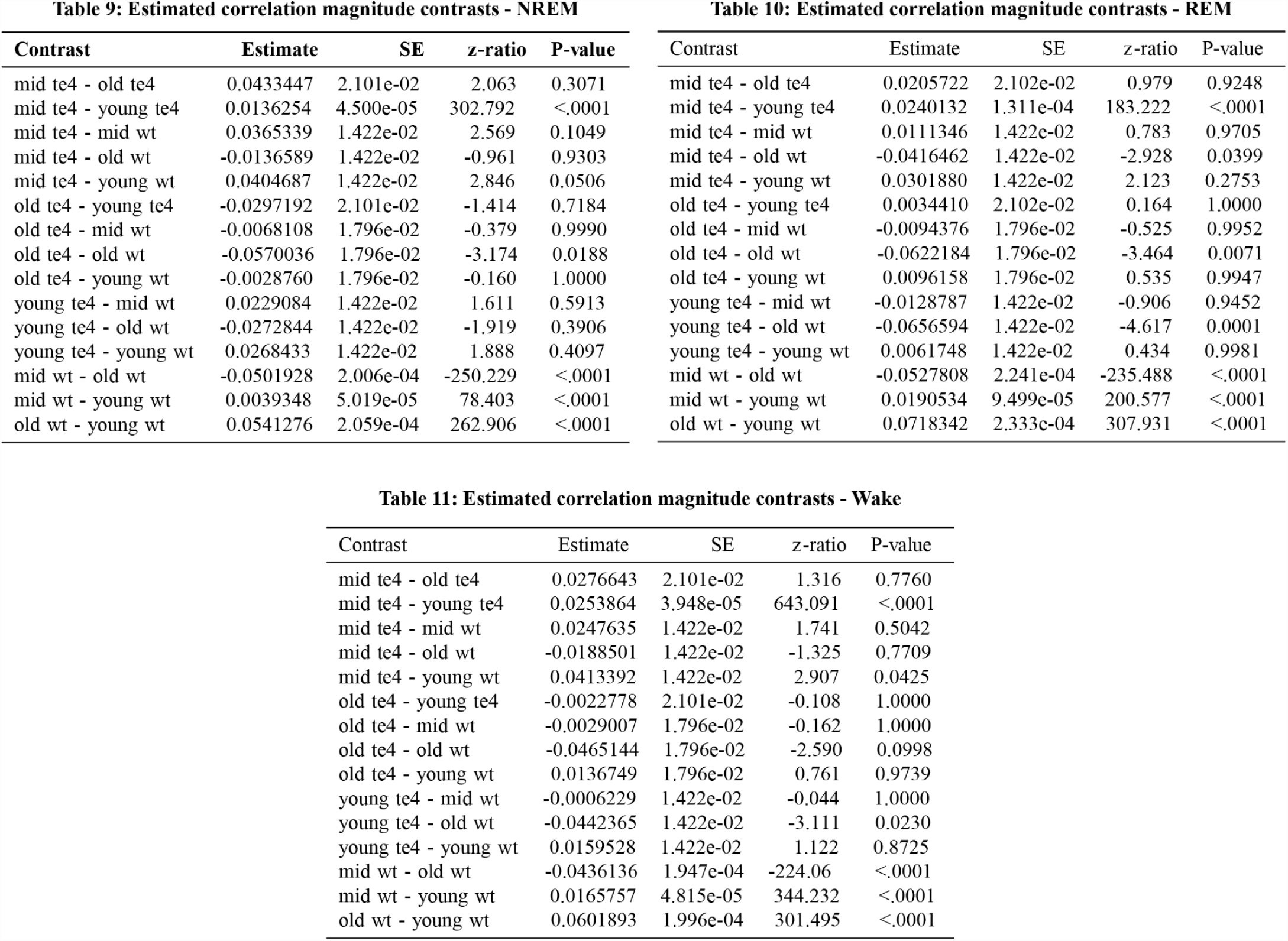
Statistical comparisons of pairwise correlation magnitude by age, genotype and brain state. The ensemble mean pairwise correlation magnitude was calculated for each brain state. The data were fit using a linear mixed effects model with the formula: corr ∼ age_group * genotype * state * cell_type + (1∣animal) where ’animal’ is included as a random effect to account for repeated measures and inherent variability across different animals. Pairwise comparisons were conducted using the emmeans package. Tables provide contrasts among the levels of the fixed effects and the associated p-values. All p-values were adjusted using Tukey’s method to account for multiple comparisons. Estimate: Predicted difference between the groups in the given contrast. SE (Standard Error): Measure of variability around the estimated difference, accounting for both fixed and random effects. z-ratio: z-ratio: The estimated difference divided by its standard error, used to test the significance of the contrast. p-value: Probability of observing the given z-ratio under the null hypothesis of no difference.

Taken together, these data suggest that there are subtle but reliable changes in correlation magnitude and complexity across the lifespan of TE4 animals, and that these changes are not a result of acute instability. Through the lenses of hierarchical bootstrapping and bootstrapped linear mixed effects regression, magnitude and complexity effects were not robustly evident (Supplemental Figure 6), underscoring their modesty. However, considered alongside the robustness of firing rate and variance, these data suggest that the impact of tauopathy on neuronal activity may be better captured in analyses of higher-order features.

### Population activity in WT CA1 hippocampus suggests a critical set-point

Neuronal activity in isocortical networks is homeostatically regulated around a distinctive nonequilibrium regime of population dynamics called “criticality” (Hsu and Beggs, 2006; Ma et al., 2019). Critical systems are precisely balanced at the transition between ordered and chaotic dynamics, a point that generates remarkable information processing capabilities (O’Byrne and Jerbi, 2022). Given the immediate relationship between near-critical dynamics and the capacity of a network to perform complex tasks (Cramer et al., 2020), a clear prediction about NDDs emerges: cognitive/behavioral disruptions caused by NDDs should directly correspond to deviations from criticality.

We first sought to determine whether spontaneous neuronal activity in WT/E4 CA1 is suggestive of a near-critical set-point. Briefly, at criticality, population events of all sizes and durations are observed: the system is scale-free. Here, population events are cascades of action potentials observed across an ensemble (Figure 5A), termed “neuronal avalanches” (Beggs and Plenz, 2003. See methods). If avalanches are scale free, their size and duration distributions must obey a power law. Then, the most stringent test of criticality dictates that the relationship between avalanche size and duration must scale according to a third, predetermined power law (Figure 5B) (Friedman et al., 2012; Tang and Bak, 1988; Touboul and Destexhe, 2017). A network’s nearness to criticality can be quantified as the difference between the predicted power law and the observed relationship, i.e., the deviation from criticality coefficient (DCC; Figure 5B). A DCC of ∼ 0.2 is expected of a network tuned to a near-critical point (Ma et al., 2019; Xu et al., 2022).

**Figure 5.**
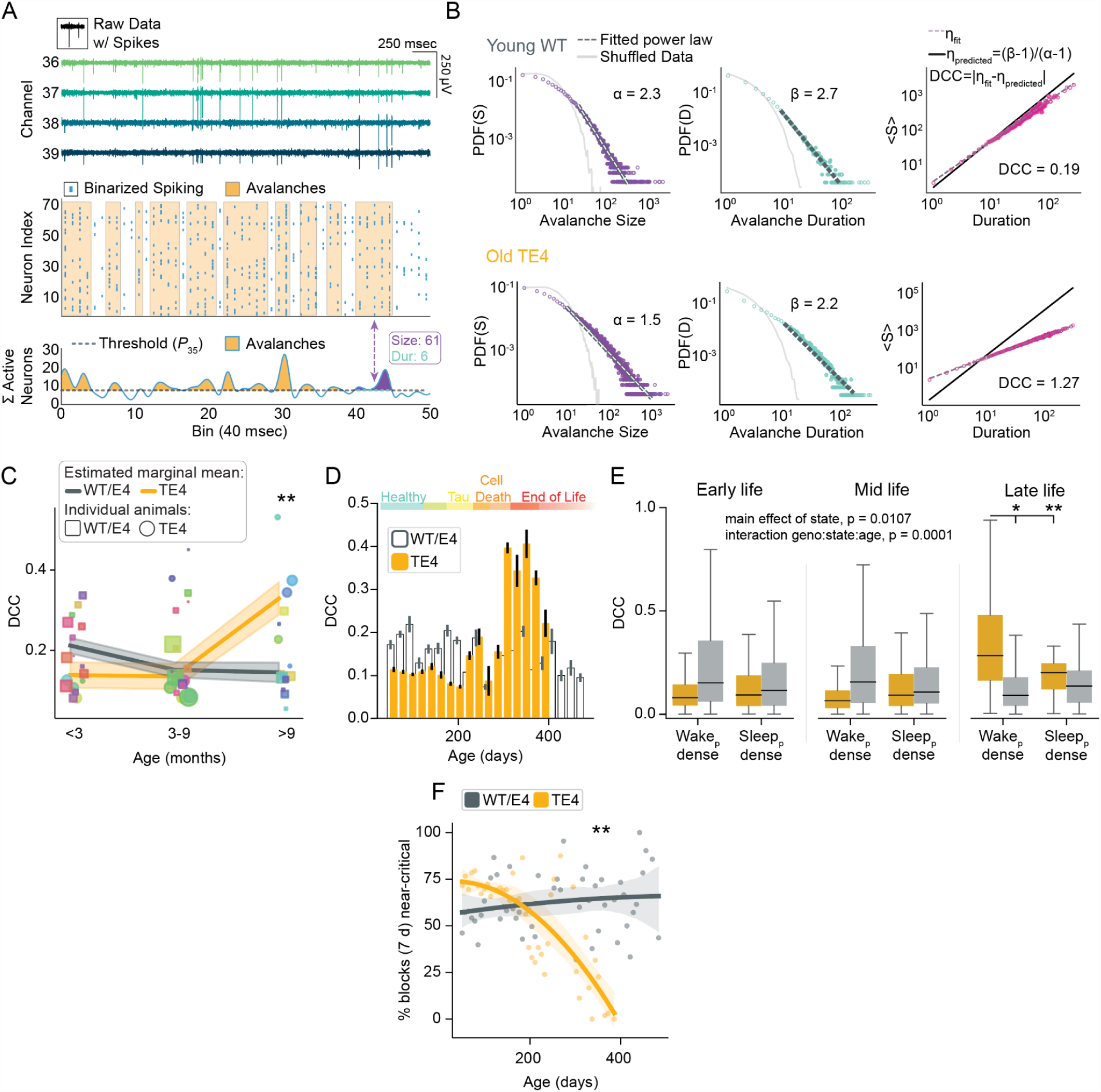
Network criticality is severely and progressively degraded by tauopathy. (A) Demonstration of neuronal avalanching. (top) Example of raw electrophysiological data from 4 channels in a recording from a WT mouse. (middle) Neural spiking was binarized in 40 ms bins and (bottom) network activity was integrated. Neuronal avalanches are defined by periods where network activity is above a threshold (dashed pink line, bottom). The size of a neural avalanche is the integrated network activity, and avalanche duration is the amount of time network activity remains above the threshold. (B) Example avalanche size (left) and duration (middle; units are number of 40 ms bins) distributions (log-log axes) fit by power laws for a WT (top) and TE4 (bottom) mouse. The scaling relation (right) between size and duration distributions shows the difference between the empirical scaling relation (dashed line fitted to pink data) and predicted scaling relation (solid line). This difference is the deviation from criticality coefficient (DCC). (C) Estimated marginal means of DCC measured in 2 h blocks were extracted from the linear mixed model that accounts for individual animals and variable durations of observation (dcc ∼ genotype * age_group + (1|animal) where animal is included as a random effect). Estimated marginal means are plotted as lines across age group and colored by genotype (error is model SE). Scatter points show the means of individual animals, indicated by color. Squares are WT/E4 animals, circles are TE4 animals. Asterisks show WT/E4 versus TE4 contrast. (D) Mean DCC values, binned in 3 week intervals, of WT and TE4 animals across the lifespan. (E) DCC as a function of genotype, age group (young, middle, and old), and brain state (wake_p_- and sleep_p_- dense: 2 h blocks of > 66% in one state). The figure displays the main effects and interactions as determined by a linear mixed model regression, formulated as DCC ∼ genotype * age_group * state_dense + (1∣animal). ’Animal’ is included as a random effect to account for within-animal correlations. Pairwise post-hoc comparisons were conducted using estimated marginal means to quantify the differences between specific group levels. (F) The proportion of data blocks across the lifetime of TE4 and WT/E4 animals that are near critical - defined as a) passing power law tests, and b) exhibiting DCC < 0.3. Asterisks denote a significant interaction of age and genotype (near_critical ∼ age * genotype + 1|animal, where animal is included as a random effect and where age is included in 7 d bins). * p < 0.05, ** p < 0.01

### Scale-free activity and the near-critical regime are disrupted by tauopathy

Examined in 4,677 intervals of 2 h, the majority of WT/E4 data were scale free and near-critical across all animals and ages (Figure 5B, top). In 3,954 intervals of 2 h, many observations of TE4 dynamics were similarly near-critical. However, in TE4 data, there was a dramatic effect with age; DCC increased progressively until eventual near-complete failure of power law structure in old TE4 (Figure 5B, bottom). There was a significant interaction of age and genotype (p = 3.165e-06, linear mixed effects: dcc ∼ age_group * genotype + 1|animal, where animal is included as a random effect), which was reflective of an increase in DCC in old TE4 animals but not WT/E4 (Figure 5C; old TE4 vs old WT p = 0.0013, linear mixed effects). The effect of age on DCC was robust to the alternate statistical approaches of a hierarchical bootstrap (old TE4 vs WT/E4 P_boot_ = 0.0092) and a bootstrapped linear mixed model (p < 0.05 in 92/100 tests of old TE4 versus old WT/E4; Supplemental Figure 7). These approaches found no differences in WT/E4 animals as a function of age.

The binning of age into young, middle, and old (Shi et al., 2017) potentially obscures the relationship between near-critical dynamics and the timecourse of disease onset. Examined in 23 bins of 3 weeks (P50 - P530), WT/E4 animals were stably near-critical (i.e., DCC <∼0.2) throughout all of life (Figure 5D). In contrast, TE4 animals were stably near-critical until approximately P200, at which point there was an inflection and DCC progressively increased to a peak in late life consistent with the end stage of disease. Given that these animals begin to accumulate insoluble tau aggregates at ∼P200 (Shi et al., 2017), this suggests that measurements of the hippocampal computational regime track progressively with the timecourse of tauopathy and may provide insight into the earliest stages of molecular pathology.

### Tauopathy disrupts the hippocampal computational regime in a state-dependent manner

There is increasing evidence that the molecular underpinnings of NDDs accumulate in a state-dependent fashion (Kang et al., 2009), such that sleep is believed to decrease (Xie et al., 2013) and wake increase their levels (Holth et al., 2019). Recent work suggests that, in healthy animals, a core function of sleep is to maintain near-critical dynamics (Xu et al., 2022). Taken together, these observations inform the hypothesis that, in NDDs, sleep may fail to reassert criticality as the disease unfolds. To address this, we examined DCC in sleep_p_- and wake_p_- dense blocks (2 h with > 66% spent asleep or awake, respectively, Supplemental Figure 3) as a function of age and genotype (Figure 5E). Consistent with recent work in the isocortex of young rats (Xu et al., 2022), DCC was slightly but significantly lower during sleep_p_ than wake_p_ in WT/E4 animals (wake_p_: 0.22 +/- 0.022, sleep_p_: 0.187 +/- 0.022; p = 0.0014, linear mixed effects). In early life, TE4 animals revealed no significant difference in DCC between sleep_p_ (0.173 +/- 0.045) and wake_p_ (0.122 +/- 0.042, p = 0.24). In mid life, TE4 sleep_p_ DCC was slightly but significantly higher than wake_p_ (0.169 +/- 0.041 versus 0.128 +/- 0.041, p = 0.0094). In old TE4 animals, the emergence of severely non-critical dynamics was exacerbated in wake_p_ (0.351 +/- 0.054 versus 0.233 +/- 0.061 in sleep_p_, p = 0.0061). These data suggest that, throughout the lifetime of TE4 animals, the effect of sleep and wake on the hippocampal computational regime may be aberrant. In old TE4 animals, the observation that DCC during sleep_p_ is close to the near-critical threshold of 0.2 implies that, while sleep may be capable of acutely achieving near-critical dynamics, the restorative effect fails to persist upon waking.

### DCC can be used to predict genotype

To test the ability of DCC to predict an animal’s genotype (Shmueli, 2010), we trained an XGBoost (Chen et al., 2016) to identify TE4 and WT animals based only on DCC (Supplemental Figure 7C). We trained on 100 randomly selected datapoints from all but two animals, withholding a WT/E4 and TE4 animal for testing. We then iterated across every pair of WT/E4 and TE4 animals to achieve an estimate of the classifier’s accuracy. Based on a single measurement of DCC, the XGBoost performed slightly above chance in young (< 3 months) animals (54 +/- 0.61%), at chance in the middle age group (3-9 months; 51 +/- 0.58%), and significantly above chance in the old animals (> 9 months; 63 +/- 1.24%). To test whether passing the model more data would improve its accuracy, we trained and tested a new set of models on multiple measurements of DCC (contiguous samples). Unsurprisingly, models that observed more samples performed better in both the young and old age groups (performance in 9-sample-point models: young = 57 +/- 1.36%, middle = 51 +/- 1.57%, old = 79 +/- 1.49%).

Average DCC may arise by one of two possible scenarios. First, the system may simply operate further from criticality as disease progresses. This would be indicative of a drifting set-point. Alternatively, impaired homeostatic processes may still be operating around a critical set-point, albeit less effectively. In this case, the system would be expected to occasionally return to criticality, but with progressively decreasing probability. To evaluate these alternate explanations, we examined the proportion of 2 h data blocks that passed power law tests as a function of age and genotype. In old WT/E4 animals, 68 +/- 5.9% of 2 h blocks of data exhibited power laws in both the size and duration distributions. In contrast, 31 +/- 13.1 % blocks passed power law tests in old TE4 animals. These data suggest that, in old TE4 animals, critical dynamics still serve as a homeostatic set-point. The dramatic increase in mean DCC is accounted for by larger, more frequent, and longer-lasting departures from the set-point as a function of disease. This is readily captured by simply evaluating the fraction of an animal’s data that satisfied the criteria of a near critical system, i.e., data a) pass power law tests (Supplemental Figure 7D) and b) exhibit a DCC of < 0.3. Measuring the percentage of data near critical shows that WT/E4 animals are stable throughout life, while TE4 animals monotonically decline from the beginning of our observations (Figure 5F).

### Disruptions in criticality correlate with molecular and anatomical markers of tauopathy

There is extensive inter-animal variability in the timing and severity of NDDs, which is recapitulated in TE4 mice (Shi et al., 2017). In other words, age is an indirect measure of disease status. While our data thus far compare age to dynamics, we reasoned that disruptions in homeostatic set-points might reliably indicate underlying pathology. Alternatively, compensatory processes (such as those at the level of firing rate) might delay the impact of disease on dynamics, rendering the relationship nonlinear. To distinguish between these possibilities, we directly compared individual circuit dynamics with molecular and anatomical markers of disease progression. TE4 animals were sacrificed intermittently between P96 and P393. We compared dynamics at the end of life with hyperphosphorylated tau (p-tau, as indicated by AT-8 immunoreactivity), hippocampal atrophy, and ventricular expansion.

There was no relationship between histological disease markers and either firing rate or CV (Figure 6A). Measures of local circuit correlations generally revealed non-significant trends that reflected disease markers (Figure 6B). The exception to this was the entropy of the pairwise correlation matrix, which varied significantly with underlying damage (p = 0.026, r^2^ = 0.48). Further, hippocampal volume, ventricular volume, and AT8 coverage each covaried significantly with the proportion of blocks near critical in the last month of each animal’s recording (Figure 6C). These data suggest that disruptions in criticality track the extent of underlying anatomical and molecular pathology.

**Figure 6.**
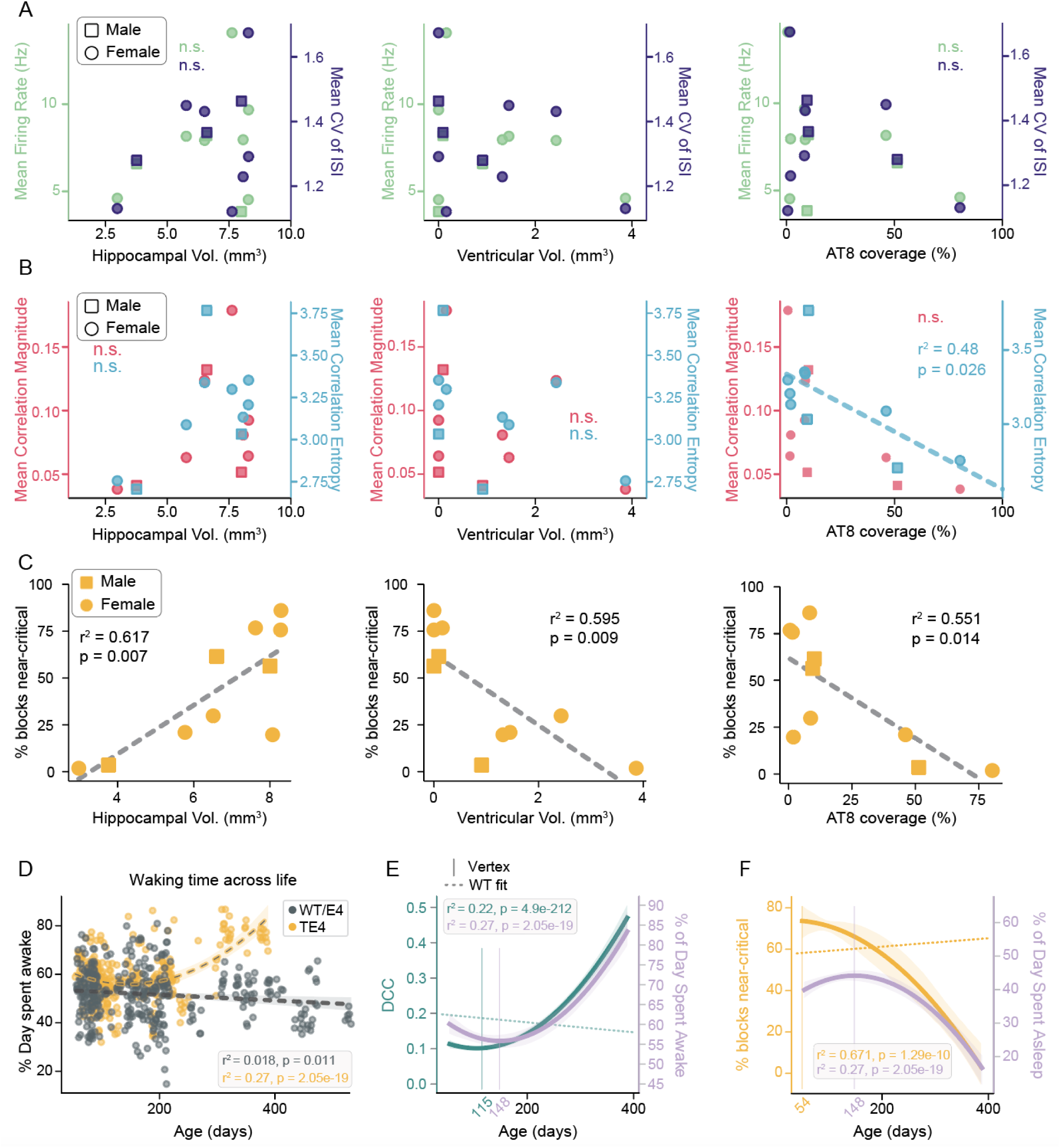
Network nearness to criticality is predictive of underlying molecular/anatomical disease markers and sleep/wake symptoms. Measures of neuronal activity in the last 1 month of each animal’s life are correlated with three measures of tau pathology in the same animals. (A) (left) Mean firing rate (light green) and CV of ISI (purple) do not significantly correlate with hippocampal volume. (middle) The same as left but for ventricular volume. (right) The same as left but for quantification of p-tau, as indicated by AT8 immunoreactivity (coverage % is the total area of the hippocampus with positive label). (B) (left) mean pairwise correlation magnitude (pink) and mean entropy of the pairwise correlation matrix (blue) do not significantly correlate with hippocampal volume or ventricular volume (middle). However, correlation entropy predicts AT8 accumulation (right). Note that, in contrast to firing rate and CV, correlation measures exhibit trends in all cases. (C) All three measures (hippocampal volume, ventricular volume, and p-tau coverage) significantly vary with the proportion of blocks that are near critical (i.e., pass power law tests *and* have DCC of < 0.3). (D) The proportion of the 24 h day spent awake_p_ is plotted against animal age for both WT/E4 and TE4 animals. Each point represents one 24 h block of a single animal. WT/E4 data are best described by a linear model, while TE4 data are best described by a second-order polynomial. (E) The best-fit model of TE4 wake_p_ over time (lavender with error) is plotted alongside the best-fit model of TE4 DCC over time (teal with error). The WT/E4 best-fit over time is shown (teal dashed line without error). The vertices of the TE4 DCC and wake_p_ models are indicated by solid vertical lines of the same color. (F) Same as D but showing percent of blocks that are near critical alongside the sleep_p_ (1 - wake_p_) over time.

### Departure from criticality tracks disrupted sleep/wake cycles

A state of disease requires symptoms (Bennett et al., 2006), however, behavioral abnormalities often lag behind pathophysiological processes (Sperling et al., 2011). To determine the relationship between disruption of criticality and symptoms, we compared DCC to alterations in sleep/wake cycles, which are some of the earliest reliable behavioral changes in NDDs (Ju et al., 2013; Lucey et al., 2021).

In TE4 but not WT mice, there was a progressive increase in the proportion of each day spent in wake_p_ (Figure 6D. Interaction of age and genotype p = 0.0065, old TE4 versus old WT/E4 p = 0.001, linear mixed effects; Holth et al., 2017). To estimate the relationship between progressive changes in sleep/wake and DCC (Figure 6D,E), we fit a series of increasing order polynomial regressions to sleep/wake measures over time and to DCC over time, and used the Akaike Information Criterion (AIC) to evaluate fit (Akaike, 1974). In WT animals, both DCC and sleep/wake measures were best fit by linear models (WT DCC: y = -0.00014x + 0.203, r^2^ = 0.008 p = 1.31e-09. WT proportion of day spent awake: y = -0.012x + 53.9, r^2^ = 0.018, p = 0.011). In TE4 animals, however, each variable was best modeled by a second-order polynomial, reflective of the non-linear, progressive nature of tauopathy (TE4 DCC: y=4.4e-06x^2^-0.001x+0.2, r^2^ = 0.22, p = 4.88e-212. TE4 proportion of day spent awake: y = 0.0005x^2^ - 0.14x + 66.35, r^2^ = 0.27, p = 2.05e-19). As a quantitative estimate of the point at which disease began to alter DCC and sleep in TE4 animals, we extracted the vertex of the best fit models by measuring where the first derivative crossed zero (Figure 6E, solid vertical lines). Dysregulation of sleep and DCC were estimated to begin at P148 and P115, respectively. Given that each measure was extracted from the same datasets, this raises the possibility that breakdown in emergent set-points may precede detectable changes in sleep structure.

Similarly, we assessed the relationship between disruptions of the sleep/wake cycle and the percentage of blocks near-critical over time (Figure 6F). The percentage near-critical for WT/E4 data was best-fit by a linear function (y = 0.04*x + 49.81, r^2^ = 0.084, p = 0.065), whereas the percentage of blocks near-critical was best modeled by a second-order polynomial in TE4 animals (y=-0.0006x^2^-0.034x+73.36, r^2^ = 0.671, p = 1.29e-10). We calculated the vertex of the best fit model of TE4 percent near critical (Figure 6F, solid vertical lines); this indicated that the proportion of data satisfying criteria to be considered near critical is continuously decreasing from P50, the very beginning of our view into tauopathy (Fig. 5F).

## DISCUSSION

It is widely understood that for neural circuits to function properly, multiple aspects of neuronal activity must be homeostatically constrained (Abbott and Nelson, 2000). Neurodegenerative disease (NDD) is noteworthy for the broad and progressive failure of circuits to function properly. Thus, we reasoned that NDD should involve the persistent and progressive deterioration of homeostatic set-points in neuronal activity. Much to our surprise, neuron-level set-points were resilient to both age and tauopathy. In contrast, a measure of the emergent network computational regime, criticality, was disrupted in parallel with the development of pathophysiology. Deviations from the critical set-point were predictive of animal genotype after the onset of pathophysiology, and correlated with anatomical and molecular markers of disease as well as the emergence of disrupted sleep cycles. Failures in critical dynamics were exacerbated in waking and either occurred with or preceded disruptions in the sleep/wake cycle, a commonly noted symptom that predicts cognitive decline in clinical populations (Lucey et al., 2019; Winer et al., 2019). These results establish that, while homeostatic mechanisms regulating single unit activity, such as firing rate, may be robust to disease, the mechanisms that stabilize information flow across networks may be highly sensitive to disease progression. Measurement of brain activity with respect to criticality thus has potential to be a powerful biomarker, and is a candidate locus connecting molecular mechanisms with symptoms.

Set-points in the activity of individual neurons appeared robust to the accumulation of intracellular tau, even in late-stage disease. This is not to suggest that neurons did not die, only that those observable by extracellular recording were indistinguishable from neurons in healthy brains. A key implication is that the mechanisms supporting firing rate homeostasis are resilient in the face of long-lasting changes in the intracellular and local circuit environment. Given that (based on first principles) disease must degrade some aspect of neural signaling, this leaves two possibilities. *First*, NDDs do not persistently disrupt neuronal activity, i.e., disruptions are restricted to acute events that were averaged out in extended recordings, or *second*, perturbation occurs at the level of emergent dynamics. Our data suggest that criticality, a feature of activity across a network, is highly sensitive to tauopathy. Retrospectively, it is perhaps not surprising that the fine tuning of complex interactions may be the node most sensitive to disease. Criticality requires the precise coordination of inhibition and excitation across large and recurrent networks. Modeling suggests that it requires plasticity of broadly connected inhibitory interneurons, thus demanding synaptic plasticity, balanced activity, and intact connectivity (Stepp et al., 2015; Poil et al., 2012). In contrast, firing rate homeostasis can occur in complete synaptic blockade and demands only that a single neuron be able to detect its own postsynaptic activity (Turrigiano et al., 1998; Ibata et al., 2008). Finally, from a purely theoretical perspective, criticality has direct explanatory power regarding the computational capacity of a network (Bak, 1996; Shew and Plenz, 2013; Cocchi et al., 2017; O’Byrne and Jerbi, 2022). As a result, any degradation of functional capacity might be expected to reflect disruption of the near-critical set-point.

On the surface, our results are incompatible with reports of NDD driving changes in neuronal firing rates. There may be a methodological explanation, as our recordings spanned tens of thousands of hours of unrestrained behavior, which provides a different view of disease than acute, more simplified preparations. However, three points merit discussion. First, both hyper and hypo activity have been suggested in tauopathy (Perez et al., 2013; Wu et al., 2021). Further, despite consistent degradation of cognition, altered activity is not observed in all NDDs. Hyperactivity is widely reported in amyloidosis (Keskin et al., 2017; Lerdkrai et al., 2018; Grienberger et al., 2012), although it may be constrained to a subset of cells (Busche et al., 2008). Second, results indicating hyper/hypo activity rely on relatively brief recordings (as little as five minutes), and are often tied to neurophysiological events and behaviors, such as sharp wave ripples (Prince et al., 2021), seizures (Palop et al., 2007), and running (Wu et al., 2021). Third, indirect measures of action potential generation, such as calcium imaging, are differentially sensitive to dissimilar patterns of neuronal activity (Wei et al., 2020; Huang et al., 2021). For example, a persistent shift from bursts towards single spikes could be mistaken as a change in rate. Together, these points raise the possibility that increased/decreased activity might be context- and disease-specific - and thus unlikely to explain persistent cognitive impairments. In contrast, it is interesting to note that, in TE4 animals > 11 months, only 9% of data were near critical. Phenomenologically, this is reminiscent of observations of fleeting moments of paradoxical lucidity in late-stage human NDD (Mashour et al., 2019; Batthyány and Greyson, 2021). While the causal relationship between the network computational regime and cognitive function remains to be established, our results clearly demonstrate that a) tauopathy can occur without measurably disrupting neuron-level set-points, and b) the network computational regime can reliably reflect disease status across long timescales.

Criticality has long been an attractive candidate for a computationally optimal set-point in the brain (Bak, 1996; Shew and Plenz, 2013; Cocchi et al., 2017; O’Byrne and Jerbi, 2022). While there is extensive evidence that brains are critical (Del Papa et al., 2017; Ma et al., 2019; Fontenele et al., 2019), the relationship of criticality to function remains largely theoretical (Hoffman and Payton, 2018; Cramer et al., 2020; but see Meisel et al., 2012). From this perspective, networks generating near-critical dynamics are assumed to be healthy and maximally capable of performing complex tasks. Combined with evidence that isocortical networks homeostatically maintain criticality in the face of perturbation (Ma et al., 2019), this supports the hypothesis that brains have evolved a suite of mechanisms to actively maintain an optimal computational regime. The possibility that NDDs should undermine criticality has been a tantalizing suggestion (Montez et al., 2009; Vyšata et al., 2014; Zimmern 2020; Habibollahi et al., 2023), although demonstration of criticality as a set-point and locus of NDDs disruption of neural function has been elusive. Here we show that hippocampal circuitry is frequently near-critical in non-pathophysiological conditions, and that criticality is uniquely sensitive to progressive tauopathy. The timecourse of the disruption of the near-critical regime in TE4 animals potentially precedes the emergence of disrupted sleep/wake cycles. While correlational in nature, these results offer strong support for the hypothesis that criticality is a key organizing principle of computationally powerful neurobiological networks. Given that the central problem in NDDs is the disruption of the ability of such networks to perform complex tasks, our data raise the intriguing question of how the restoration of criticality might impact functional decline in NDD.

The role of sleep in NDDs, particularly Alzheimer’s Disease, is noted in both human and animal studies. Sleep deprivation, disrupted sleep architecture, and irregular sleep are all linked to negative outcomes in NDDs. One plausible explanation of this lies in the state-dependent flushing of extracellular fluids, i.e., the glymphatic system (Xie et al., 2013). However, this cannot account for the increased release of intracellular tau by sleep (Holth et al., 2019; Wang and Holtzman, 2020). Another avenue by which sleep may counteract disease centers on the role of neuronal activity in determining disease progression. Aberrant activity facilitates disease (Yamamoto et al., 2015), and blockade of activity mitigates disease-impact of sleep deprivation (Holth et al., 2019). Recent work in healthy animals suggests that a central, restorative function of sleep is the resurrection of criticality (Xu et al., 2022). This suggests a possible link between critical dynamics and the rate of disease progression. Our data reveal two points consistent with this: a) wake was enriched in non-critical dynamics in TE4 animals, despite a lack of firing rate dysregulation, and b) the onset of disrupted criticality appeared to precede or coincide with early sleep disruptions (although this is based on the vertices of best-fit models, and is thus only an estimate).

We note that, even in old TE4 animals, hippocampal networks *are* near critical during sleep. As a result, we raise two possible mechanisms by which sleep disruptions may contribute to disrupted brain activity in tauopathy. The first is analogous to a car that won’t stay in gear after the shifter is released. While sleep can acutely restore a critical regime, these effects fail to persist upon waking. The second explanation is simply that animals fail to sleep enough. In this case, waking proceeds past a typical threshold and induces activity-dependent disease processes. The key difference between these explanations is that the former cannot be rescued by sleep augmentation.

Our data cannot resolve whether firing rate-independent deviations from criticality are causative in disease timing and severity. Assigning causality in complex systems is challenging, but conservatively, increased DCC and/or disruptions in power law signatures are effective biomarkers that may offer direct insight into the decline of circuit function. Further underscoring the possibility of application, estimates of criticality can be achieved non-invasively (Jannesari et al., 2020; Fosque et al., 2022). We note the paradoxical trend for young TE4 animals to be slightly but consistently closer to criticality than age-matched WT animals. This effect was strong enough that the XGBoost performed slightly above chance when classifying withheld young animals. The most parsimonious explanation for this is a simple side effect of the transgene, as TE4 animals have elevated soluble tau in early life (Shi et al., 2017). However, there is a priori reason to believe that operating too near to criticality could be detrimental. This line of argument asserts that brains in a slightly subcritical regime maintain a buffer against the possibility of super-criticality (Priesemann et al., 2014), i.e., a state of runaway gain (Meisel et al., 2012). Exogenously driven activity that exacerbates NDD (Yamamoto et al., 2015) is almost certainly biasing the system towards supercriticality, raising the possibility that NDDs may be driven by cumulative impact of these deviations from the near-critical set-point. Future work will be required to evaluate whether too-close-to-criticality is a predictive factor in other models of NDD.

## ACKNOWLEDGMENTS

This work is supported by: NIH BRAIN Initiative 1R01NS118442-01 and the Brightfocus Foundation Standard Award Program in Alzheimer’s Disease Research 29225 (KBH); NIH RF1AG047644, NIH RF1NS090934, and the JPB Foundation (DMH); NIH 1R01EB029852, NSF IIS-2146072, and the CIFAR Azrieli Global Scholars Program (ELD).

## METHODS

### Animals

The generation of TE4 animals (P301S tau/ApoE4 Sullivan ApoE KI line) is described in detail in Shi et al., 2017 and Wang et al., 2021. Briefly, human ApoE4 KI mice (C57BL/6) were crossed with P301S tau transgenic mice (The Jackson Laboratory, 008169) on a C57Bl/6 background. The P301S mouse harbors the T34 isoform of microtubule-associated protein tau with one N-terminal insert and four microtubule binding repeats (1N4R). This encodes the human P301S mutation which is driven by the mouse prion protein promoter. These mice generate tau aggregates in the context of apoE4, the most potent genetic risk factor for sporadic Alzheimer’s Disease. Note that the P301S/E4 mouse does not express pathogenic amyloid. WT and TE4 littermates of each sex were selected for recordings. The sex of animals is noted in corresponding figure legends.

All procedures were conducted in accordance with protocols approved by the Washington University in St Louis Institutional Animal Care and Use Committee (IACUC), following NIH guidelines for the care and use of research animals. In total, 25 female and 18 male mice were used (12 female and 11 male C57BL/6 mice, 5 female and 2 male E4 mice, and 8 female and 5 male TE4 mice). Mice were postnatal day 36 (P36) to P533 during recordings. All mice were housed in an enriched environment with a 12:12 h light/dark cycle and had ad libitum access to food and water during the recordings.

### Surgery

Prior to the start of surgery, adult mice were administered Buprenorphine SR (0.1 mg/kg), Dexamethasone (0.5 mg/kg) and Meloxicam (5 mg/kg) for pain relief and were provided with Ringer’s solution for hydration. Mice were anesthetized with isoflurane (4-5% for induction and 1-2% for maintenance, mixed with air). Mice were head-fixed in a robotic stereotaxic instrument and the fur, skin, and periosteum covering the dorsal surface of the skull was removed. A craniotomy 1 - 1.5 mm in diameter was cut above CA1 hippocampus and dura was resected. A custom-built, 64-channel, tetrode-based multielectrode array was inserted into CA1 (-2.54 mm AP and +/-1.75 mm ML relative to bregma, -1.5 mm DV relative to brain surface) at a rate of 5mm/minute using a robot and a vacuum-based probe holder. A ground/reference wire was connected to a screw implanted anterior on the skull, and the multielectrode array was connected to headstage electronics (WhiteMatter, LLC), enclosed in a 3D printed case, and secured with dental cement. Mice were administered meloxicam (5 mg/kg) and dexamethasone (0.5 mg/kg) for 3 d of recovery prior to the start of neural recordings.

### In vivo recording and spike sorting

Neural data were recorded continuously from freely behaving animals in the context of the home cage for up to 200 d. Electrical signals were amplified, sampled (25 kHz), and digitized on the head of the animal and sent via cable to an eCube server system (WhiteMatter). Video of the mice was recorded at 15 or 30 frames per second, synchronized with electrophysiological data.

In parallel to ongoing recordings, raw data from the prior day were divided into 2, 4, 8, or 12 h intervals for identification of units based on compute availability (number of detected units, unit firing rates, DCC, etc. were unaffected by block size). Blocks were first bandpass filtered (500 to 7,500 Hz) and waveforms that crossed a -4 standard deviation threshold were extracted for clustering. Single units were identified using a modified version of MountainSort4 and XGBoost classifier (key parameters learned by the classifier include waveform shape, waveform consistency, interspike interval distribution features, multi-channel pickup, presence ratio, signal-to-noise etc). Single units were manually confirmed by experienced humans blinded to condition. In this work, we included units labeled ‘Quality 1’ and ‘Quality 2’, where ‘Quality 3’ is multi-unit and ‘Quality 4’ is contaminated by noise. We recorded a total of 179,478 Quality 1 and 2 single-units across all animals and ages. On average, each individual clustering job returned 40.46 +/- 5.74 single-units from the 64 channel array.

### Estimation of Cell Type

Well-isolated single-units (Supplemental Figure 1) were separated into putative cell classes based properties established by prior work on cell types in CA1 (Skaggs et al., 1996; Csicsvari et al., 1999; Frank et al., 2001; Mizuseki et al., 2009, 2012). Cells were first broken into putative fast spiking inhibitory interneurons (pFS) and regular spiking units (RSU; putative excitatory) based on peak latency (time from the peak of the waveform to the trough). Neurons were classified as putative fast-spiking (pFS) if peak latency was < 0.3 ms. Amongst the RSU population (peak latence > 0.3 ms), principal neurons are identified as those whose mean rate across the entire clustering interval was < 5 Hz (Skaggs et al., 1996; Csicsvari et al., 1999; Frank et al., 2001; Mizuseki et al., 2009, 2012). Neurons were classified as ‘other RSU’ if peak latency was > 0.3 ms but overall firing rate was > 5 Hz. For all analyses, only cells with a presence ratio >= 0.99 were used (spiking activity was present during 99% of the duration of the clustering block).

### Histology

At the end of each recording, mice were perfused with 4% formaldehyde (PFA). Brains were extracted and fixed in PFA for 24 h then moved to a PBS solution for an additional 24 - 48 h. Brains were sectioned at 50 μm and slices were mounted on slides. Nissl staining was performed using cresyl violet, and stained sections were compared with the Allen Institute Mouse Brain Atlas (Allen Institute for Brain Science, 2012) to confirm electrode placement (Supplemental Figure 1C).

At the end of recordings from TE4 mice, the above procedures were followed for the brain hemisphere where electrodes were surgically implanted. The other hemisphere was used to quantify markers of tauopathy, using previously established methods (Shi et al., 2017; Yanamandra et al., 2015). This hemisphere was also sliced into 50 μm sections then washed in Tris-buffered saline (TBS) buffer 3 times, and incubated in 0.3% hydrogen peroxide in TBS for 10 min. TBS washes were performed three times, then sections were blocked with 3% milk in 0.25% TBS-X (Triton X-100) for 0.5 h followed by incubation at 4°C overnight with biotinylated AT8 antibody (Thermo Scientific, MN1020B, 1:500). The next day, slices were washed three times in TBS and then incubated with the VECTASTAIN Elite ABC HRP solution (Vector laboratories, PK-6100) for 1 h to amplify the AT8 signal. Finally, sections were washed 3 times in TBS and developed using DAB solution (Sigma, D5905). Images of stained slices were quantified using Image J.

Of these brain sections, every sixth section starting rostrally at bregma -1.3 mm to the dorsal end of the hippocampus at bregma −3.1 mm were used for hippocampal volume analysis (6 – 7 sections per mouse depending on the severity of brain atrophy). These sections were stained with 0.1% Sudan black in 70% ethanol for 20 min, then washed in 70% ethanol for 1 min. This process was repeated three times, then sections were washed in Milli-Q water three times for 10 min each, then coverslipped with Fluoromount. Imaging of stained sections was performed, then the hippocampus and lateral ventricle were manually traced and quantified using NDP viewer. The volumes of these regions were calculated using the formula: volume = (sum of area) * 0.3 mm. For hippocampus and posterior lateral ventricle, quantification started from bregma −1.3 mm and ended at bregma −3.1 mm.

### Manual sleep scoring

Raw neural data was downsampled to 500 Hz from 2 - 5 channels for each recording, selected for the presence of robust single unit activity and clean local field potential (LFP). The extracted LFP (0.1– 60 Hz) was plotted on a spectrogram for each hour of the recording. The corresponding video file was analyzed using DeepLabCut (Mathis et al., 2018) for markerless pose estimation, from which movement traces were measured and synchronized with neural data. Accuracy of machine-vision based motor output was compared with EMG-based methods to confirm redundancy (see Parks et al., 2023). Sleep scoring was performed using the combination of LFP spectral power and movement: periods of time with high delta band power and low movement were labeled as NREM sleep, periods with low delta band power, high theta band power, and no movement were labeled as REM sleep, and periods with high theta and movement were labeled as wake. All labels were made in 4 s epochs. A total of 60 days (1440 h) from 30 distinct animals were manually scored: 29 days from TE4, 26 days from WT, and 5 days from E4 mice. Scored days were from animals 56 days old to 532 days old.

### XGBoost Sleep Scoring Model

A XGBoost model was trained on 20 d (480 h) of manually scored data from 12 unique animals: 9 d from TE4, 11 d from WT/E4 mice. 3,510 features from the LFP signal were extracted and used to train the model, and performance was evaluated by comparing the model predictions for 72 h of test data from 3 withheld mice (i.e., the model’s training data did not include any samples from the test animals). The balanced accuracy of the model on withheld test animals was 82.7%. This XGBoost classifier was then used to automatically score 26,729 h of the neural recording dataset from 42 out of the 43 individual animals. To evaluate the accuracy of automated scoring, we conducted several analyses to compare model predictions with human-scored sleep-wake data and established patterns recognized in the field. First, we compared the distribution of LFP theta power during NREM, REM, and wake bouts. In both the entire human-scored dataset and the entire classifier-scored dataset, the distribution of LFP theta power was highest during REM and lowest during NREM, in congruence with well-established norms. Also, in both the entire human-scored dataset and the entire classifier-scored dataset, the distribution of LFP delta power was highest during NREM and lowest during REM, again replicating well established patterns. To assess whether the classifier-scored dataset reproduced the impact of tauopathy on the sleep-wake cycle of TE4 mice found in the human-scored dataset, we grouped the sleep predictions for each animal into consecutive 24 h windows and only used windows containing a complete 24 h of predictions. We measured the percent of time spent awake within each 24 h window, then used Cook’s distance to detect highly influential data points (likely outliers) and filter them from the dataset. We found nearly identical trends in the classifier-scored dataset as the human-scored dataset: as WT mice age, the percent of each day they spend awake remains stable, whereas as TE4 mice age, the percent of each day they spend awake dramatically increases. We grouped the dataset into age ranges relevant to disease progression (< 3 months, 3-9 months, and > 9 months old, based on Shi et al., 2017), to compare the mean time spent awake in TE4 mice vs WT/E4 mice across these age ranges. Finally, again to confirm the accuracy of the classifier predictions, we measured time spent awake in all animals during the light cycle and dark cycle. In both genotypes, animals spent significantly more time awake when the lights were off than when the lights were on. This result was similar in both the classifier- and human-scored datasets and recapitulates known circadian patterns in mice.

### Firing rate analysis

Ensembles of neurons were considered in 2 h windows based on the shortest clustering block length (see above). Mean firing rate in Hz was calculated as total n spikes / 7200 s. While clustering blocks represented continuous time series, we did not track individual units across processing jobs. Linear mixed effects modeling was performed using the formula: log_firing_rate ∼ age * genotype * cell_type + 1|animal, where animal is included as a random effect. To measure state-specific set-points in firing rate, we extracted the number of spikes fired by each neuron in each epoch of REM, NREM, and wake that occurred in each 2 h block. Those counts were concatenated and the mean rate was calculated as n spikes / total time in a state. Epochs > 20 s were considered for these analyses. Linear mixed effects followed the form of log_firing_rate ∼ age * genotype * state * cell_type + 1|animal, where animal is included as a random effect.

### CV of ISI analysis

The coefficient of variation (CV) of the interspike interval (ISI) distribution was calculated across all single units using methods that have been described previously (Taube, 2010; Hengen et al., 2013). On a neuron-by-neuron basis, we considered all ISIs in 2 h windows, filtering out extreme values (ISIs greater than the mean + 4 SD), and calculated the CV (standard deviation / mean). Linear mixed effects modeling was performed using the formula: CV ∼ age * genotype * cell_type + 1|animal, where animal is included as a random effect. Mean CV of ISI values across an entire ensemble of single units was calculated by finding the CV of each individual neuron, then averaging all neurons together in each 2 h window. To calculate CV within different behavioral states, we extracted all ISI values for a given neuron across all wake bouts, all NREM bouts, and all REM bouts that were > 20 s and occurred during the time period when that neuron was recorded and clustered, then calculated the CV of by state. Linear mixed effects modeling for this dataset was performed using the formula: CV ∼ age * genotype * state * cell_type + 1|animal, where animal is included as a random effect.

### Pairwise correlation analysis

To assess pairwise correlations of neural activity from the two genotypes over age, we followed previously published methods that were used to indicate a homeostatic set-point in pairwise correlation strength (Wu et al., 2020). Spiking counts from each single-unit were measured in 100 ms bins. The Pearson correlation between the spike-counts of each pair of neurons was measured in 30 min intervals, producing a pairwise correlation matrix. We performed hierarchical clustering on the first 30 min window, then stepped this window forward in 5 minute increments across the entire length of the recording while maintaining the original row/column structure for each neuron. Autocorrelations were removed. Pairwise correlation magnitude was calculated by measuring the average absolute value of all unique pairwise correlations in the matrix. Correlation complexity was measured by finding the average entropy (bits) per pair.

To measure correlation magnitude and complexity by behavioral state, we binned single-unit spiking in 100 ms and measured the correlation between each pair of neurons, but we calculated the pairwise correlation matrix in 4 s intervals to align with the corresponding behavioral state (sleep state was labeled in 4 s intervals). We measured mean correlation magnitude and entropy across all pairwise correlation matrices arising in bouts of wake, NREM, and REM that were > 20 s and contained > 10 neurons, where a bout is defined by consecutive 4 s intervals of the same behavioral state.

To measure correlation stability, we used blocks where single-units were clustered in intervals >= 4 h (5114 total blocks, 2521 TE4). In this context, we measured the drift in correlation matrices over 4 h intervals. We binned spiking in 100 ms and created two pairwise correlation matrices: one from the first 30 min of the block and one from the last 30 min. We used hierarchical clustering from the first block and maintained the ordering of neurons in the second. We computed the L1 distance between the two correlation matrices, then normalized this value by dividing by the number of neurons. We repeated this for all 4 h intervals of time within each recording.

To assess the impact of sleep-wake states on correlation stability, we measured the pairwise correlation matrix from the first 10 min and last 10 min of each 1 h block that was scored by the sleep scoring model and measured the L1 distance between these two matrices, normalized by number of neurons. For each 1 h block, we measured the proportion of time spent awake or asleep in the first and last 10 minutes as well as the proportion of time spent awake or asleep during the entire 1 h. We defined sleep-dense as 1 h blocks were the proportion of wake during the entire 1 h and during each of the 10 minutes when the pairwise correlation matrices were calculated were < 33%. We defined wake-dense as 1 h blocks were the proportion of wake during the entire 1 h and during each of the 10 minutes when the pairwise correlation matrices were calculated were > 66%.

### Neuronal avalanches analysis and measurement of criticality

Avalanche-based analyses were performed as described previously (Ma et al., 2019; Xu et al., 2022). Spike times of each neuron were binarized in 40 ms bins, where a 0 represents no spiking activity for that neuron during those 40 ms and a 1 represents spiking activity during the window. This process was performed for every single-unit, and then the overall neuronal network activity was defined by the sum of all binarized activity across all neurons within each 40 ms time bin. A threshold was set at the 35th percentile of this integrated network activity, and neuronal avalanches were defined as successive time bins where the network activity was above this threshold. Size of a neuronal avalanche is the integrated network activity above the threshold during one avalanche, and the duration of a neural avalanche is the total number of time bins that the avalanche lasted. The size and duration of all neuronal avalanches in 2 h windows were calculated for each recording. For each 2 h window, we fit the distribution of avalanche sizes and the distribution of avalanche durations separately into truncated power-laws using maximum likelihood estimation, which produced a power-law exponent α for the size distribution and a power-law exponent β for the duration distribution (Clauset et al., 2009). The goodness of fit for each power-law distribution was evaluated by simulating 1000 artificial power law distributions with the same exponent, number of avalanches, minimum avalanche size, and maximum avalanche size as the actual avalanche distribution (Clauset et al., 2009; Ma et al., 2019). The deviation between the simulated distributions and a perfect power-law was quantified with a KS statistical test and compared to the KS value for the comparison between the actual distribution and its power-law fit. The p-value for goodness of fit was calculated as the fraction of artificial distributions with KS values smaller than the KS value of real avalanche distribution. Using a significance level of 0.05, if p < 0.05, the hypothesis that the real avalanche distribution was well-fit by the power-law was rejected, whereas for p ≥ 0.05 the power-law hypothesis was not rejected (Clauset et al., 2009; Ma et al., 2019). The scaling relation between the average size of neuronal avalanches and the corresponding duration could be modeled by the equation: ⟨S⟩(T) ∼ T^η, where η could either be fit by linear regression from experimental data or predicted based on the exponent relationship equation: η_predicted=(β-1)/(α-1). The absolute value of the difference between the predicted η and the fitted η is the Deviation from Criticality Coefficient (DCC, Ma et al., 2019; Xu et al., 2022; Habibollahi et al., 2023). Any 2 h block where the avalanche size distribution and duration distribution both passed p-value tests for power-law fits and DCC was < 0.3 was classified as “near-critical.”

To analyze DCC in wake-dense or sleep-dense blocks for each animal, we calculated the amount of wake_p_ and sleep_p_ (NREM_p_ or REM_p_) predicted by the sleep scoring model in the 2 h time window used to calculate DCC. Wake-dense blocks were defined by 2 h windows where the animal spent > 66.7 % of the time awake, and sleep-dense blocks were defined by 2 h windows where the animal spent > 66.7 % of the time asleep.

We trained XGBoost models on DCC data to predict the genotype of individual animals. To evaluate model performance, we employed a modified form of Leave-p-Out Cross Validation with class stratification. For each age group, we trained an XGBoost model on 100 samples (resampled if necessary) of DCC data from each animal except for one WT/E4 mouse and one TE4 mouse, which were intentionally withheld from the training dataset. We then tested the model on 100 samples from each of these two withheld mice and measured the proportion of correctly classified genotypes. We then repeated this process for each possible combination of withheld animals, generating an array of balanced accuracy scores for that age group and DCC feature size. Balanced accuracy is reported as mean +/- SEM unless otherwise indicated. We observed little improvement on the training set as a function of hyperparameter tuning, and therefore elected to use the default values rather than tuning them on a held-out validation set (Chen et al., 2016). To test whether the inclusion of increasing amounts of DCC data improves model performance, we systematically performed this training/testing procedure: first on 100 samples containing 2 contiguous DCC points, then on samples with 3 contiguous points, followed by 4, and so on, continuing up to samples with 9 contiguous DCC points. Model performance was assessed at each stage based on the number of contiguous features included.

### Linear mixed effects regression

Statistical analyses were performed using linear mixed-effects models to assess the influence of age, genotype, cell type, and behavioral state, as well as their interactions on the dependent variable(s). Analyses were conducted using the lme4 package in R (Bates et al., 2015). In these models, "animal" was incorporated as a random effect to account for within-subject variability. The model formula was specified as follows: Response Variable ∼ Age + Genotype + Cell Type + Behavioral State + (1|Animal).

Following the construction of the linear mixed-effects models, estimated marginal means were computed to assess the mean response at each level of the fixed factors, while accounting for random effects. Pairwise comparisons among the estimated marginal means were subsequently carried out using Tukey’s honest significant difference (HSD) test. Analyses of estimated marginal means were performed using the emmeans package in R (Searle et al., 1980; github.com/rvlenth/emmeans). Standard error (SE) in emmeans is calculated as the square root of the variance of a linear predictor.

### Hierarchical bootstrap test

To further address the hierarchical structure of our dataset (within genotype there are animals, and within animals there are neurons, etc), we used a hierarchical bootstrap (Saravanan et al., 2020; McGregor et al., 2022) to compare conditions. As an example, to analyze mean firing rate by genotype and age group, we generated 10,000 bootstrapped means per condition. To do so, we randomly resampled at each level of hierarchy in our dataset, and repeated this process 10,000 times for each genotype: each sample was a randomly selected animal and within that a randomly selected neuron, whose mean firing rate was calculated. To assess statistically significant differences between genotypes, we calculated a joint probability distribution for the means of the resampled datasets and measured the percentage of this distribution that was above the unity line. This percentage is P_boot_. Using an acceptable type 1 error rate of 0.05, any value of this P_boot_ ratio greater than 0.975 indicates the mean of dataset 1 was significantly greater than the mean of dataset 2, and any value less than 0.025 indicates the mean of dataset 1 was significantly less than the mean of dataset 2. P_boot_ values between 0.025 and 0.975 indicate no significant difference between the two datasets.

Error is calculated as standard deviation. When employing the hierarchical bootstrap method, standard deviation of the resampled means is equivalent to the standard error of the original dataset (Saravanan et al., 2020).

### Bootstrapped linear mixed effects models

To assess whether a statistically significant difference between conditions would be detectable in more traditionally constrained experimental approaches, we used linear mixed models on bootstrapped datasets. Specifically, we randomly selected three to twelve 2 h bins of data from each animal and constructed a linear mixed model to test whether conditions differed using emmeans as described above. We repeated this process 100 times to generate a range of p-values expected from acute observations of neuronal activity in our datasets.

---

**Supplemental Figure 1.**
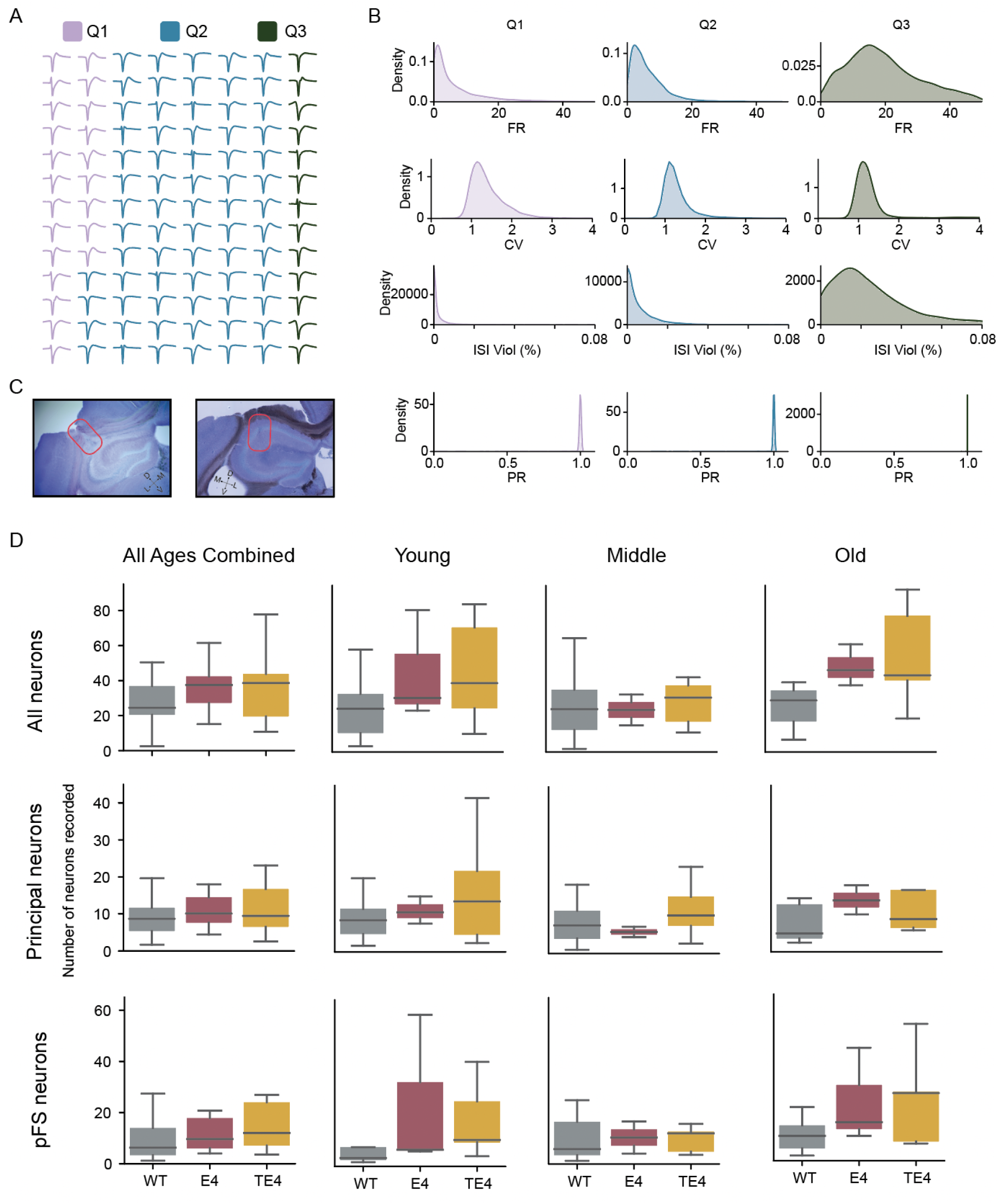
Continuous, long-term recordings exhibit high-quality single-unit activity. Example data shown from an old TE4 animal > 1 month after implantation. (A) This illustrates the mean waveform from each spike-sorted unit. The color indicates the quality of each unit. Q1 and Q2 are single units, while Q3 represents multiunit clusters that were excluded from this work. (B) This panel showcases histograms of the distribution of various unit quality metrics across all units shown in A. In order: mean firing rate, coefficient of variation (CV), ISI violations (the percentage of ISIs violating the minimum 1 ms refractory period), and presence ratio (PR), a measure of the fraction of time in which a unit was observable. (C) Example histology revealing electrode tracts that terminate in the medial/lateral extent of CA1. (D) Box plot of the mean number of single units observed in a clustering block as a function of age, genotype, and cell type (64 channel tetrode-based arrays implanted chronically without drives).

**Supplemental Figure 2.**
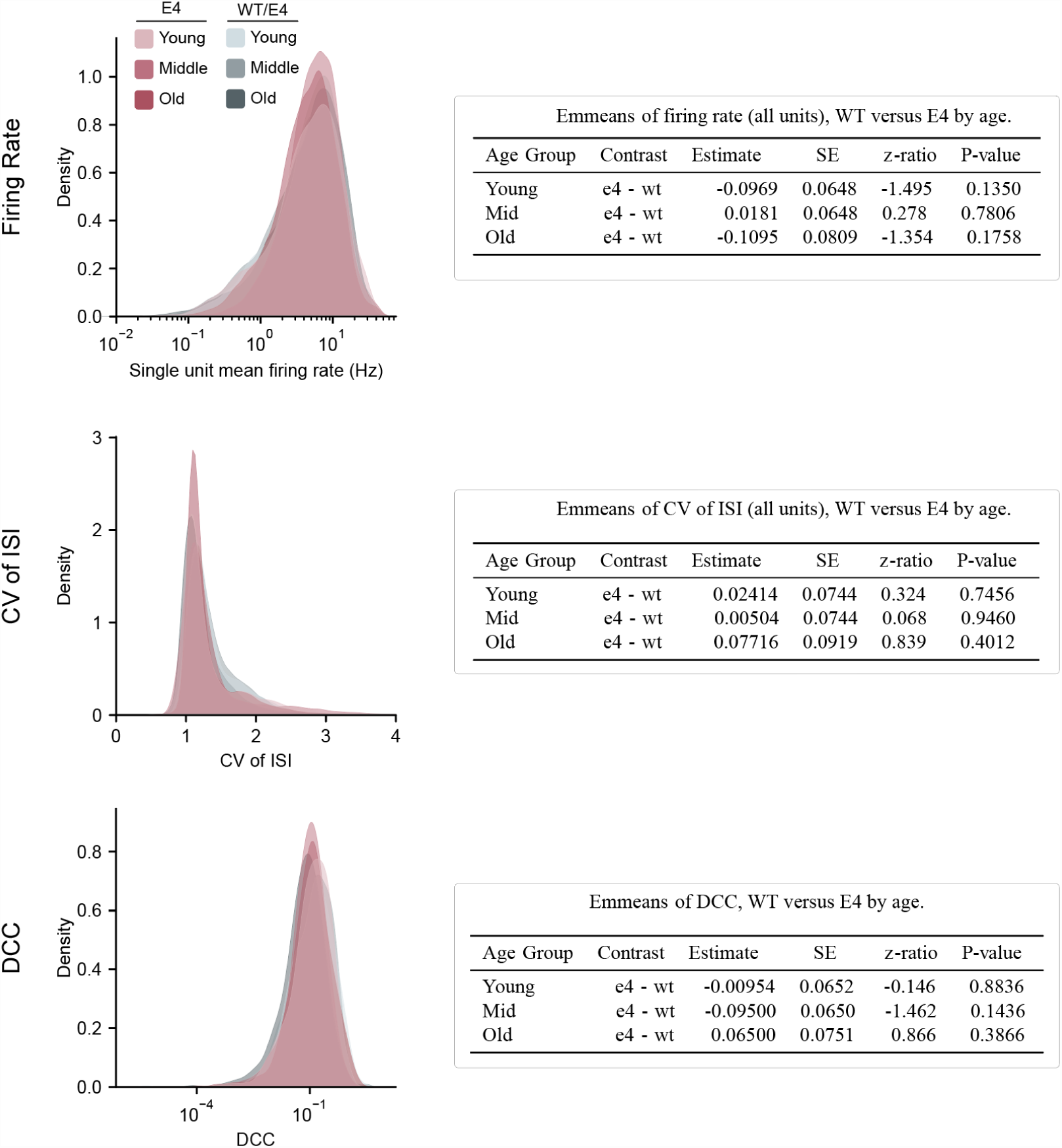
Homeostatically regulated features of spontaneous neuronal activity are not significantly different between WT and E4 animals at any age. WT and E4 animals were compared as a function of single unit mean firing rate (top), CV of ISI (middle), and the deviation from criticality coefficient (DCC). Statistical contrasts between genotypes were evaluated in three age categories: young, middle, and old. No comparisons revealed a statistically significant p value. Linear mixed effects: log_firing_rate ∼ genotype * age + (1|animal), cv ∼ genotype * age + (1|animal), and dcc ∼ genotype * age + (1|animal), where animal is included as a random effect. Pairwise comparisons were conducted using the emmeans package. Tables provide contrasts among the levels of the fixed effects and the associated p-values. All p-values were adjusted using Tukey’s method to account for multiple comparisons. Estimate: Predicted difference between the groups in the given contrast. SE (Standard Error): Measure of variability around the estimated difference, accounting for both fixed and random effects. z-ratio: z-ratio: The estimated difference divided by its standard error, used to test the significance of the contrast. p-value: Probability of observing the given z-ratio under the null hypothesis of no difference.

**Supplemental Figure 3.**
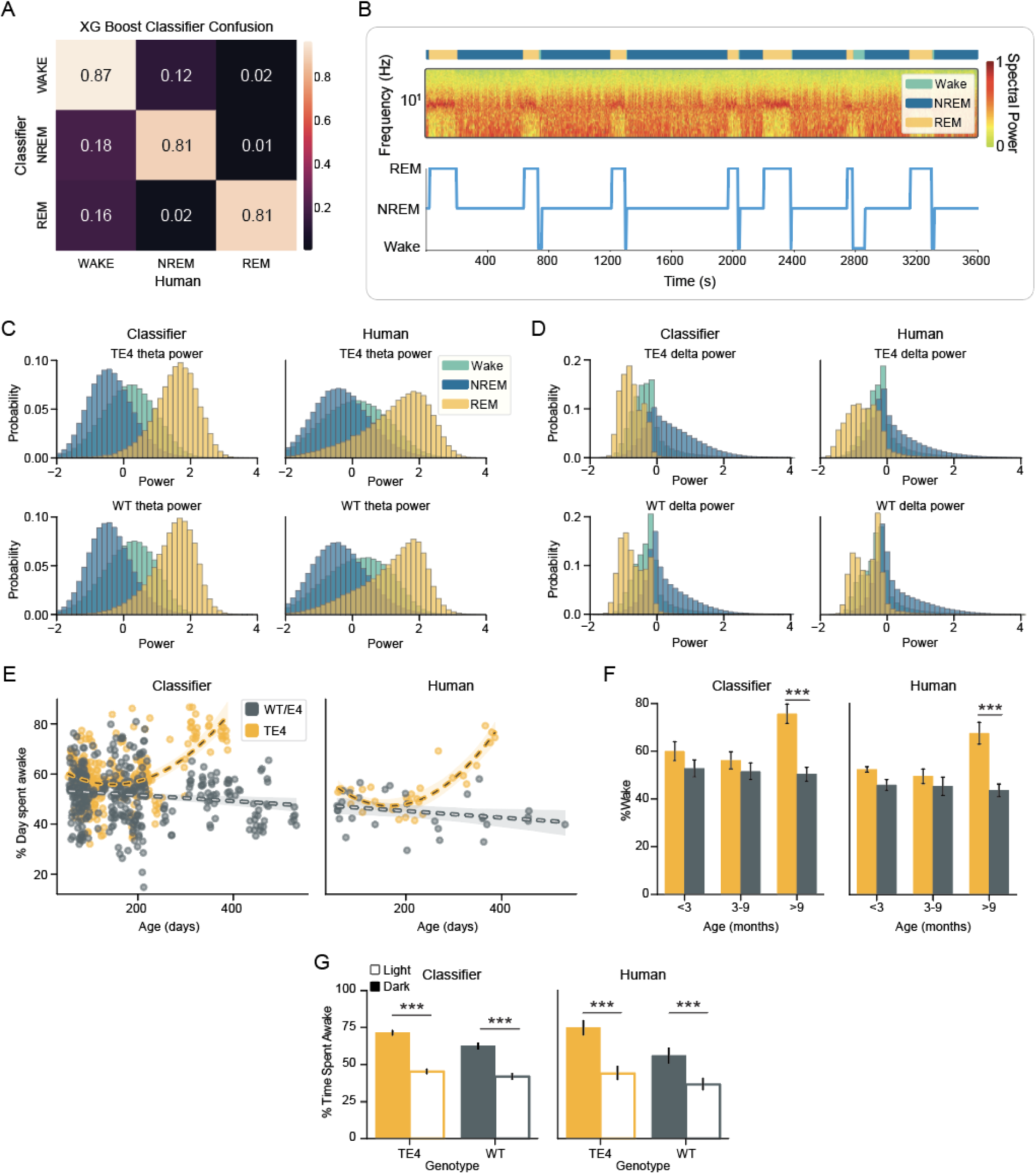
Validation of fully-automated sleep scoring by an XGBoost classifier trained and tested only on neural data. (A) Confusion matrix describing the performance of the XGBoost model when tested on three withheld animals (i.e., not in the training dataset) that were hand-scored by human experts for comparison. (B) Example output from the XGBoost. Top area plot shows model-based state labels above the relevant spectrogram (0.1-60 Hz) calculated from the input data (middle). The bottom shows the XGBoost labels in a traditional hypnogram. Note the microarousals that consistently occur following REM bouts. (C) Probability distributions of theta power (6-8 Hz) by state in XGBoost and manually labeled data in each of WT and TE4 datasets. (D) The same as C but for delta power (0.1-4 Hz). (E) The proportion of a 24 h day spent awake as a function of time and genotype, calculated based on XGBoost labeling (left) and manual labeling (right). (F) Quantification of data in E. (G) The proportion of time spent awake during the light cycle versus the dark cycle, as a function of genotype (WT/E4 and TE4) when labeled by the classifier (left) and by human (right). * p < 0.05

**Supplemental Figure 4.**
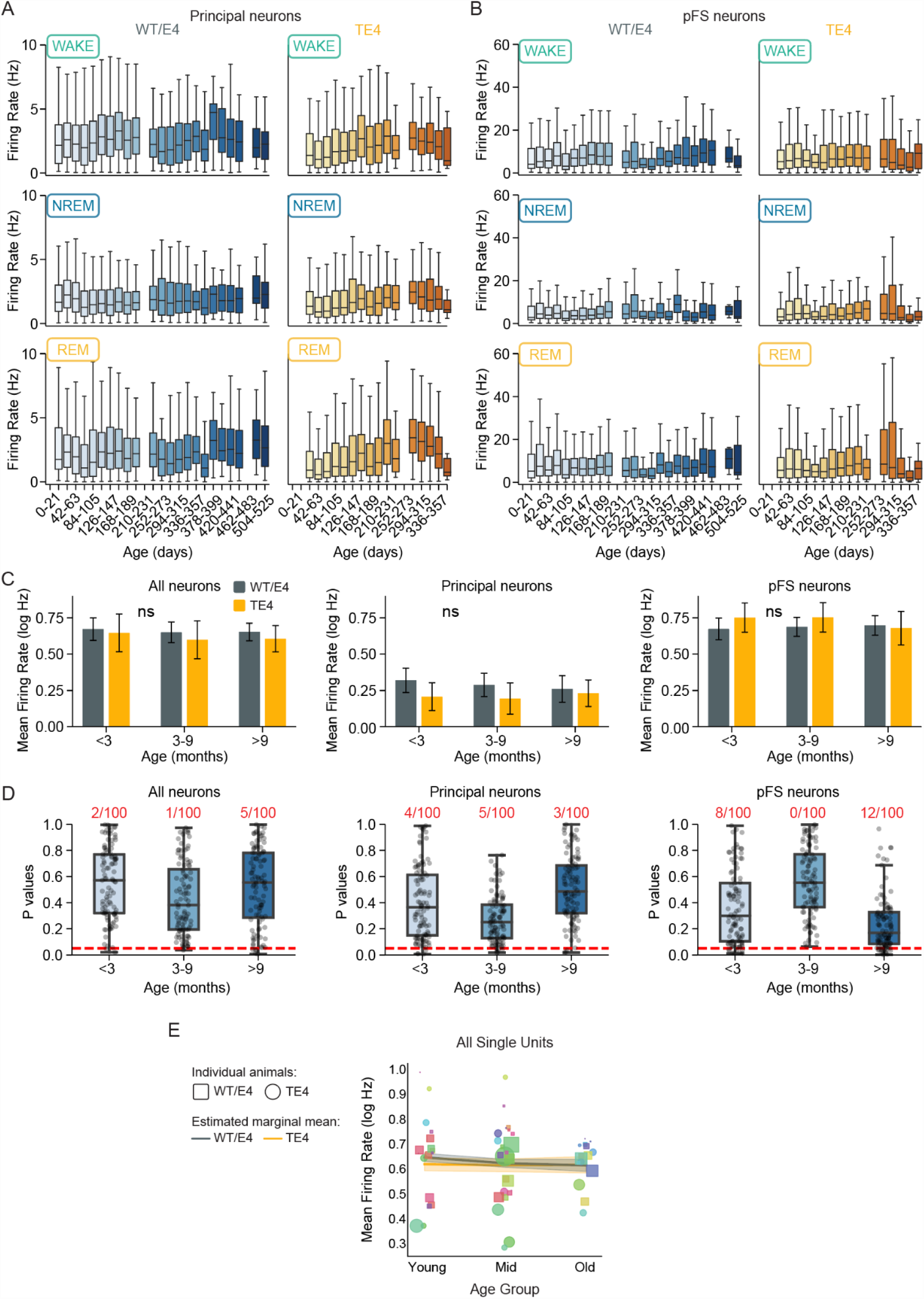
Single unit firing rates as a function of cell type, genotype, and age. (A) Mean firing rate of principal neurons across the lifetime as a function of brain state (rows) and genotype (columns). (B) Same as A but for pFS neurons. (C) Results of hierarchical bootstrap statistical tests comparing firing rates of all neurons (left), principal neurons (middle) and pFS neurons (right) as a function of genotype and age. No significant differences were observed. (D) Results of bootstrapped linear mixed effects regression comparing mean log firing rate as a function of cell type as in C and age (logfiring_rate ∼ genotype * age_group * cell_type + (1|animal) where animal is included as a random effect). The number of tests that produced a p < 0.05 is printed above each bar in red. (E) Estimated marginal means of firing rates of all single units were extracted from the linear mixed model that accounts for individual animals and variable durations of observation. Estimated marginal means are plotted as lines across age group and colored by genotype (error is model SE). Scatter points show the means of individual animals, indicated by color. Squares are WT/E4 animals, circles are TE4 animals.

**Supplemental Figure 5.**
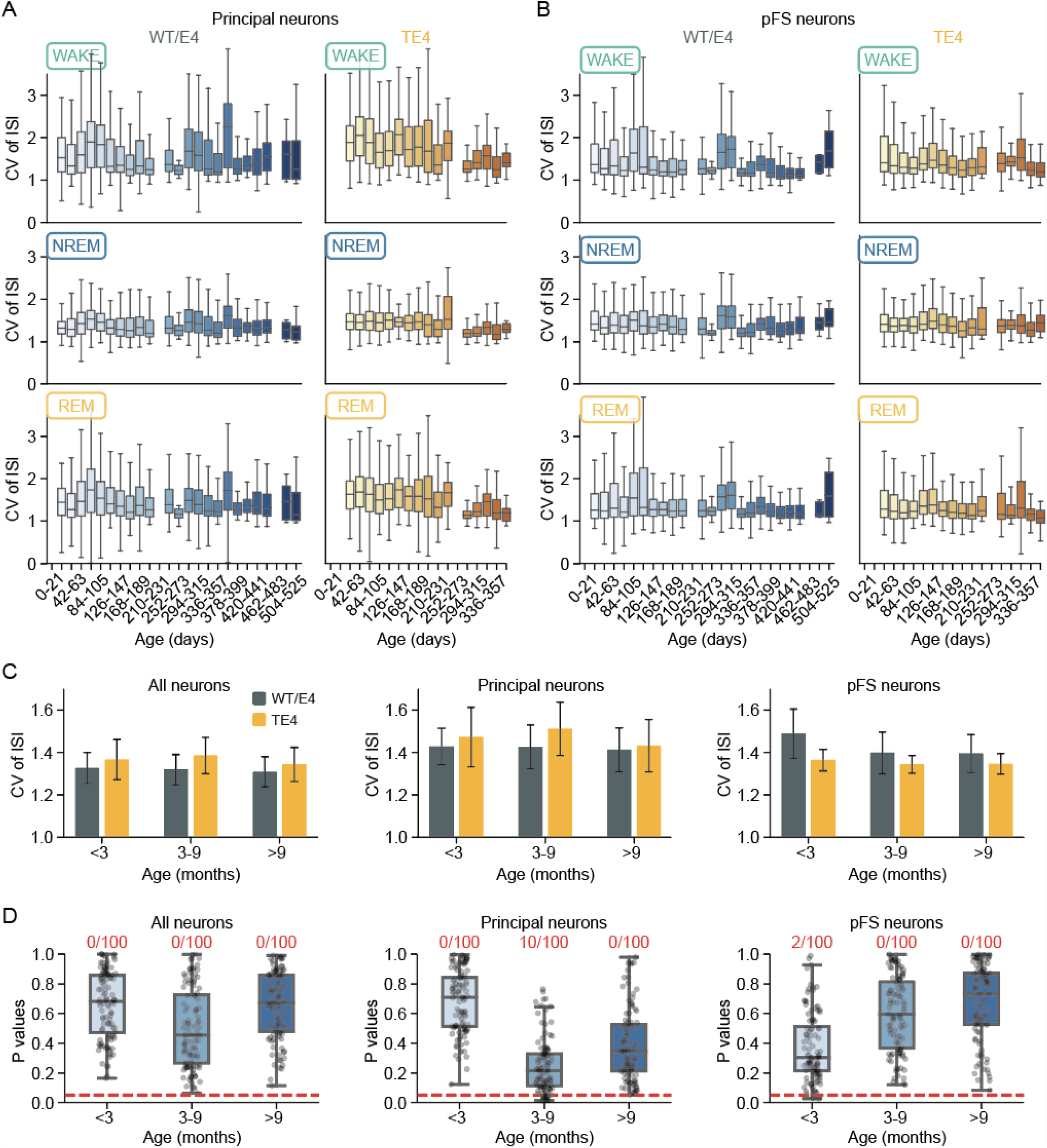
Single unit ISI CV as a function of cell type, genotype, and age. (A) Mean ISI CV of principal neurons across the lifetime as a function of brain state (rows) and genotype (columns). (B) Same as A but for pFS neurons. (C) Results of hierarchical bootstrap statistical tests comparing CV of all neurons (left), principal neurons (middle) and pFS neurons (right) as a function of genotype and age. No significant differences were observed. (D) Results of bootstrapped linear mixed effects regression comparing mean CV as a function of cell type as in C and age (cv ∼ genotype * age_group * cell_type + (1|animal) where animal is included as a random effect). The number of tests that produced a p < 0.05 is printed above each bar in red.

**Supplemental Figure 6.**
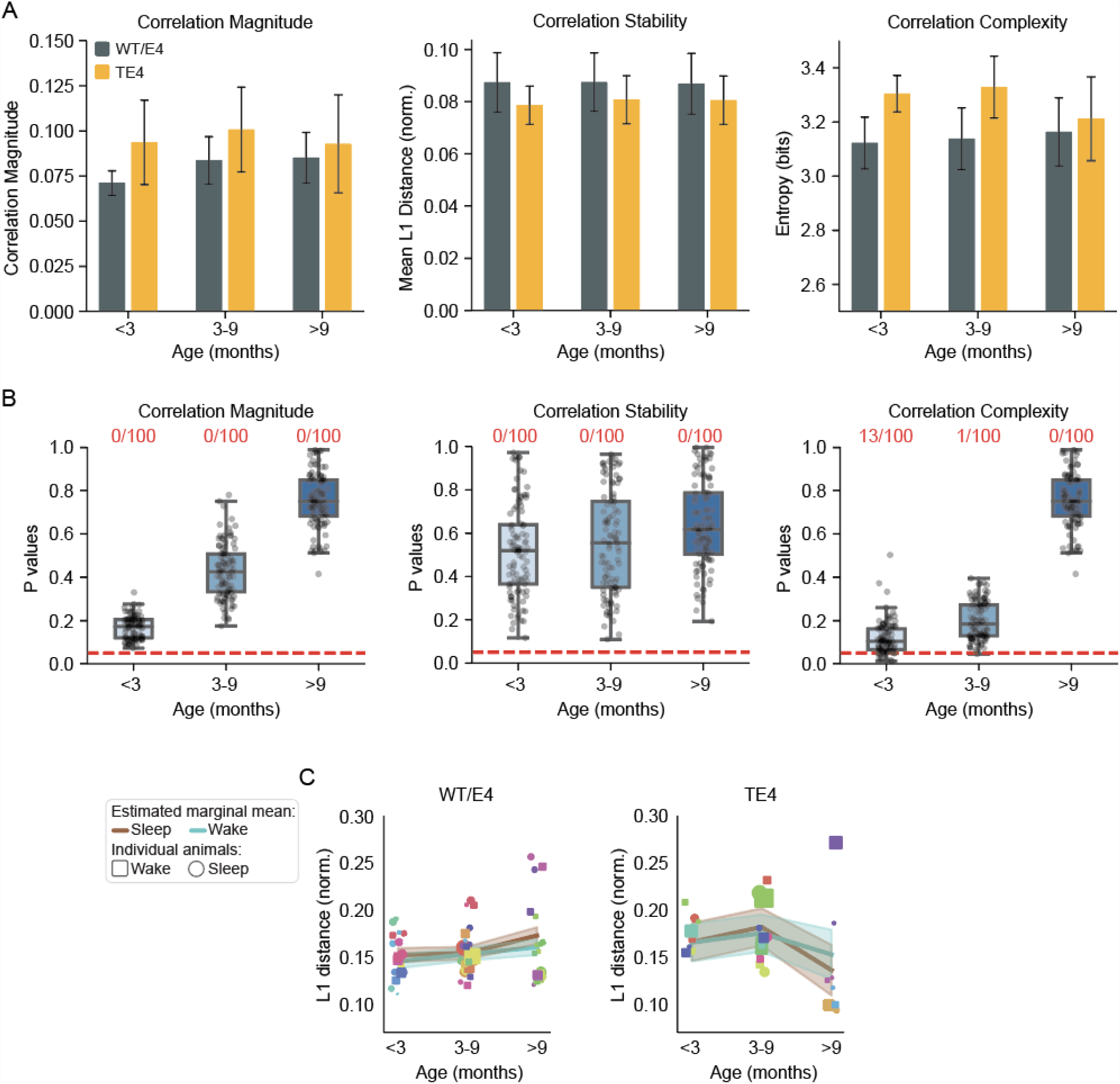
Pairwise correlation supplemental statistics. Three features of all pairwise correlations in recorded ensembles were evaluated: the mean pairwise correlation magnitude, the entropy of the pairwise correlation matrix (i.e., complexity), and the stability of the matrix. (A) Results of hierarchical bootstrap statistical tests comparing magnitude (left), stability (middle), and complexity (right) as a function of genotype and age. No significant differences were observed. (B) Results of bootstrapped linear mixed effects regression comparing correlation magnitude (left), stability (middle), and complexity (right) as a function genotype and age (corr_feature ∼ genotype * age_group + (1|animal) where animal is included as a random effect). The number of tests that produced a p < 0.05 is printed above each bar in red. (C) Estimated marginal means of normalized L1 distance - a measure of matrix change - as a function of sleep_p_-dense and wake_p_-dense blocks (1 h) across young (< 3 months), middle (3 - 9 months), and old (> 9 months) ages. WT/E4 is plotted on the left, and TE4 on the right. Estimated means were extracted from the linear mixed model that accounts for individual animals and variable durations of observation. Estimated marginal means are plotted as lines across age groups and colored by sleep_p_/wake_p_ (error is model SE). Scatter points show the means of individual animals, indicated by color. Squares are wake_p_-dense, circles are sleep_p_-dense.

**Supplemental Figure 7.**
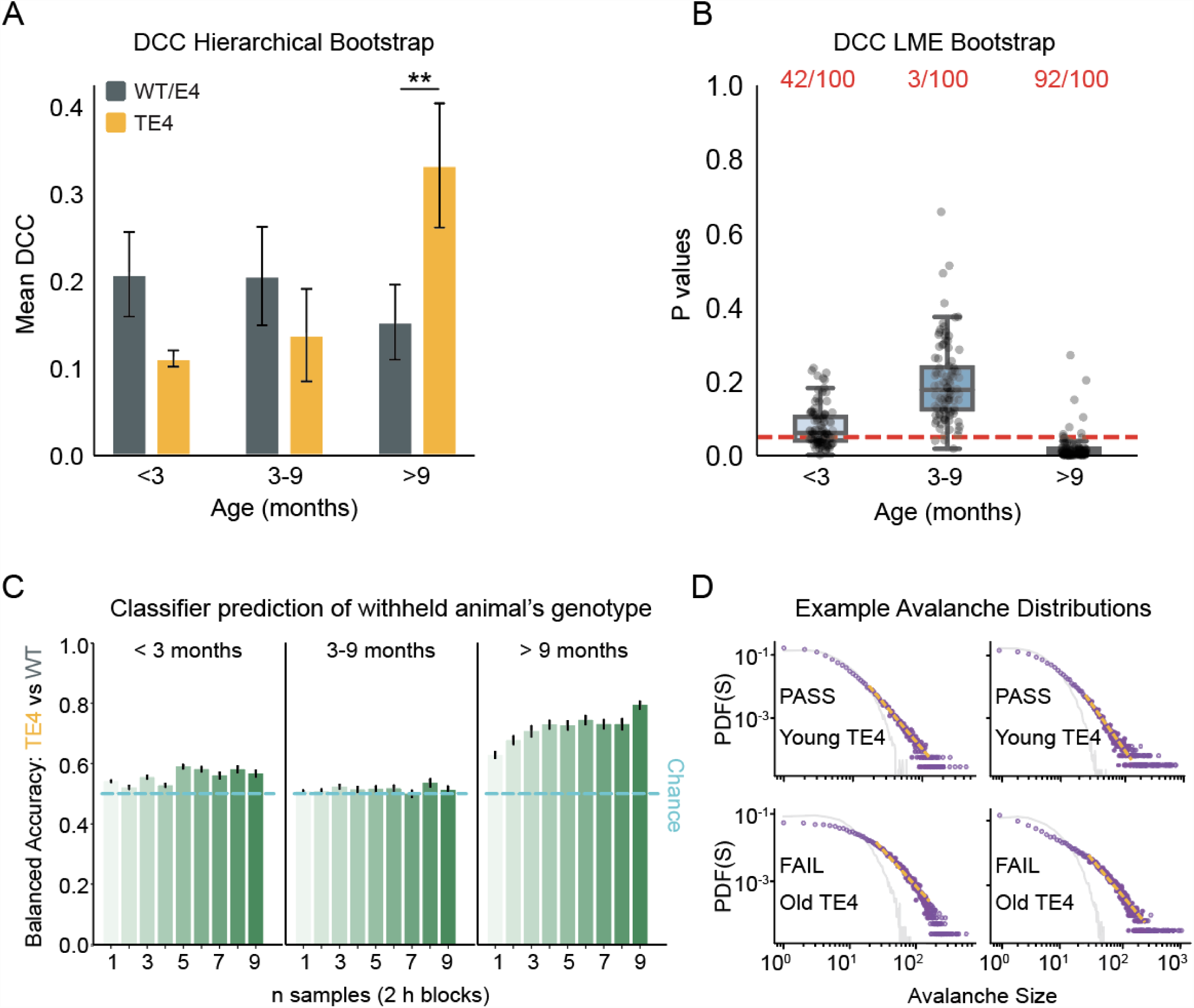
DCC statistical tests, genotype prediction, and example distributions. (A) Results of hierarchical bootstrap statistical tests comparing DCC as a function of genotype and age. This revealed a significant difference between WT/E4 and TE4 animals older than 9 months (old TE4 mean DCC = 0.333 +/- 0.071 vs old WT/E4 mean DCC = 0.153 +/- 0.043, P_boot_ = 0.0092) (B) Results of bootstrapped linear mixed effects regression comparing DCC from TE4 vs WT/E4 mice in different age groups. (C) Balanced accuracy of an XGBoost classifier trained to predict genotype (WT/E4 vs TE4) of withheld animals based on observations of DCC. Models were trained and tested on samples of 1 to 9 temporally adjacent DCC values. Chance is exactly 50%. (D) Four example avalanche size distributions. The top two pass powerlaw tests, while the bottom two fail power law tests.

